# NAD^+^ capping of RNA in Archaea and Mycobacteria

**DOI:** 10.1101/2021.12.14.472595

**Authors:** Olatz Ruiz-Larrabeiti, Roberto Benoni, Viacheslav Zemlianski, Nikola Hanišáková, Marek Schwarz, Barbora Brezovská, Barbora Benoni, Jarmila Hnilicová, Vladimir R. Kaberdin, Hana Cahová, Monika Vítězová, Martin Převorovský, Libor Krásný

## Abstract

Chemical modifications of RNA affect essential properties of transcripts, such as their translation, localization and stability. 5’-end RNA capping with the ubiquitous redox cofactor nicotinamide adenine dinucleotide (NAD^+^) has been discovered in organisms ranging from bacteria to mammals. However, the hypothesis that NAD^+^ capping might be universal in all domains of life has not been proven yet, as information on this RNA modification is missing for Archaea. Likewise, this RNA modification has not been studied in the clinically important *Mycobacterium* genus. Here, we demonstrate that NAD^+^ capping occurs in the archaeal and mycobacterial model organisms *Methanosarcina barkeri* and *Mycobacterium smegmatis.* Moreover, we identify the NAD^+^-capped transcripts in *M. smegmatis,* showing that this modification is more prevalent in stationary phase, and revealing that mycobacterial NAD^+^-capped transcripts include non-coding small RNAs, such as Ms1. Furthermore, we show that mycobacterial RNA polymerase incorporates NAD^+^ into RNA, and that the genes of NAD^+^-capped transcripts are preceded by promoter elements compatible with σ^A^/σ^F^ dependent expression. Taken together, our findings demonstrate that NAD^+^ capping exists in the archaeal domain of life, suggesting that it is universal to all living organisms, and define the NAD^+^-capped RNA landscape in mycobacteria, providing a basis for its future exploration.

## INTRODUCTION

Nicotinamide adenine dinucleotide (NAD^+^) is a universal cofactor present in all living organisms and necessary for energy metabolism, DNA repair, as well as for other critical processes (1). In 2009 it was found that *Escherichia coli* and *Streptomyces venezuelae* contain RNA with covalently attached NAD^+^ (2), thereby suggesting the presence of transcripts with 5’-NAD^+^ in both gram-negative and grampositive bacteria. Soon afterwards, the identity of NAD^+^-modified transcripts was determined in *E. coli* (3). Since then, the presence of the 5’-NAD^+^ modification and the identity of NAD^+^-RNAs have been revealed in *Bacillus subtilis* (4), *Staphylococcus aureus* (5), yeast (6, 7), plants (8–10), mitochondria (6, 11, 12), and human cells (13, 14). Moreover, Bird *et al.* (2016) demonstrated that the attachment of NAD^+^ to RNA is carried out co-transcriptionally (15). Namely, RNA polymerase (RNAP) can incorporate NAD^+^ as the first dinucleotide during transcription initiation at promoters that contain A:T (non-template: template strand) at the transcription start site (TSS, or +1) (15). The mechanism of non-canonical transcription initiation with NAD^+^ was termed NAD^+^ capping (16).

The universal, vital role of NAD^+^, together with the finding of 5’-NAD^+^ modified RNAs in prokaryotic and eukaryotic organisms led to the idea that 5’-NAD^+^ capping of RNA may be universal in all three kingdoms of life (i.e., Bacteria, Eukarya and Archaea) (17, 18) but so far NAD^+^ capping has only been detected in Bacteria and Eukarya. Hence, the first aim of this study was to analyse archaeal RNA from *Methanosarcina barkeri* to search for NAD^+^ caps. *M. barkeri* is an anaerobic methanogenic microorganism that ferments various carbon compounds and is involved in a number of biotechnological and ecological processes (19, 20); there is even a debate about the potential biological origin of Martian methane by similar organisms (21). The results of our analysis demonstrate that 5’-NAD^+^-RNAs exist in Archaea, and, therefore, NAD^+^ capping is a universal RNA modification, common to all domains of life.

Next, given the high diversity of gram-positive bacteria, we investigated the occurrence of 5’-NAD^+^ in Mycobacterium, a clinically important genus of Actinobacteria. This genus includes serious pathogens such as *Mycobacterium tuberculosis* and *Mycobacterium lepreae*, which continue to threaten public health, especially in developing countries (www.who.int). *Mycobacterium smegmatis* is another member of the *Mycobacterium* genus, widely used as a model organism to study mycobacteria. *M. smegmatis* is a non-pathogenic, fast-growing bacterium that shares a high percentage of orthologous proteins and physiological characteristics with *M. tuberculosis* (22). Therefore, we studied the composition of the NAD^+^-capped transcriptome in *M. smegmatis* in different growth phases, and we identified the NAD^+^-capped transcripts.

In summary, this work demonstrates the existence of NAD^+^ caps in Archaea and mycobacteria, and provides the first identification of the NAD^+^-capped transcripts in Actinobacteria, paving the way for future studies of NAD^+^ capping in these taxa.

## MATERIALS AND METHODS

### Strains and growth conditions

*M. smegmatis* strain mc^2^155 was grown as previously described (23). Briefly, cells were streaked from glycerol stocks onto 7H10 agar medium plates supplemented with glycerol (0.5 % v/v) and cultured at 37 °C for 2 days. Then, the agar-grown cells were inoculated into 20 ml of 7H9 medium supplemented with 0.2 % glycerol and 0.05 % Tween 80 and grown aerobically overnight at 37 °C. This overnight preculture was used to inoculate 1 L of the same medium to an initial OD_600_ ≈ 0.01. The culture was grown until OD_600_ ≈ 0.5 (exponential phase) or 24 hours (stationary phase), and cells were harvested by centrifugation (11,325 g, 10 minutes, 4 °C), and the cell pellets were immediately frozen and stored at −80 °C. *Methanosarcina barkeri* was cultivated in modified Widdel medium (1 g/L NaCl; 0.4 g/L MgCl_2_·6H_2_O; 0.15 g/L CaCl_2_·2H_2_O; 0.5 g/L KCl; 0.2 g/L KH_2_PO_4_; 0.25 g/L NH4Cl; 1 ml/L trace elements SL10, (24) 1 ml/L selenite-tungstate solution (25); 30 ml 1 M NaHCO_3_). The medium was stored in glass bottles secured with rubber stoppers and perforated caps. Air was replaced with H_2_/CO_2_ (4:1 v/v). After sterilization, the medium was supplemented with the following compounds per liter: 0.25 g Na_2_S·H_2_O; 0.087 g L-cysteine; 0.05 mg cyanocobalamin; 0.15 g pyridoxin-dihydrochloride; 0.1 mg nicotinic acid; 0.05 g calcium-D(^+^)-pantothenate; 0.04 mg 4-parabenzoic acid; 0.01 mg D(^+^)-biotin,; 0.05 mg riboflavin; 0.1 mg thiamine hydrochloride. Finally, acetate and methanol were added to a final concentration of 30 mM, and pH was adjusted to 7.0. The media were inoculated by the addition of 5 % of 1-week-old preculture, and the gas was replenished to an absolute pressure of 1.8 Bar. The cells were incubated at 36 °C for one month, with periodical additions of acetate and methanol, to a final OD_578_ ≈ 0.25.

### RNA isolation

For NAD captureSeq, total RNA from *M. smegmatis* was extracted following the protocol described in (26) with minor modifications. Briefly, the cells from 0.5 L culture were collected by centrifugation at 11,325 g, 10 minutes at 4 °C. The pellet was resuspended in 2.4 ml of TE 1X with 0.6 ml of LETS buffer (0.1 M LiCl, 0.01 M EDTA, 0.01 M Tris-HCl (pH 7.4), 0.2 % SDS), followed by the addition of 3 ml of acidic phenol. The cell suspension was sonicated on ice, three times for 1 minute with 1 minute intervals, using a ultrasonic processor set at 1 cycle and 50 % amplitude (UP200S, Hielscher). The aqueous phase was sequentially extracted with equal volumes of acidic phenol:chloroform (1:1) and chloroform, then mixed with 0.1 volumes of 3 M sodium acetate and an equal volume of isopropanol, and incubated at −20 °C overnight. The RNA was precipitated by centrifugation (24,560 g, 30 minutes at 4 °C), and the pellet was washed with 75 % ethanol and air-dried for 5 minutes. Finally, the RNA pellet was resuspended in ultrapure water (Braun) and treated with Turbo DNase (Invitrogen) according to the manufacturer’s instructions. The RNA used for other purposes (Ms1 pull-down, enzymatic treatment) was extracted similarly, with a starting volume of culture of 5 ml. For this, the volumes of TE and LETS buffers and acidic phenol were escalated down 10 fold, and a recovery marker was added before the addition of acidic phenol (24 ng of m^7^G-RNAI or 4 ng of m^7^G-A-less controls). For RNA extraction from *M. barkeri,* approximately 2 L of culture were divided into 50 ml falcon tubes and centrifuged at 7,690 g, 20 minutes at 4 °C. Pellets were resuspended in 2.5 ml of 1 X TE supplemented with 0.5 mg/ml lysozyme and 2.5 ml of Lysis Buffer (200 mM NaCl, 20 mM Tris-HCl pH 7.5, 40 mM EDTA, 0.5 % SDS), and the samples were treated with proteinase K (1 g/L) at 37 °C for 30-60 minutes. Then, an equal volume of acidic phenol was added, and the cells were sonicated on ice, three times for 30 seconds with 1 minute intervals. The aqueous phase was further sequentially extracted and RNA was precipitated and DNase treated as described for *M. smegmatis*.

### NAD captureSeq

To identify the NAD^+^-capped transcripts in *M. smegmatis,* we performed NAD captureSeq on biological triplicates of RNA samples from *M. smegmatis* cells in exponential and stationary phases of growth, following the protocol described in Winz *et al.,* 2017 (27). Each RNA replicate was divided into two aliquots of 20 μg of total RNA, one treated with ADP-ribosylcyclase (ADPRC, ADPRC+ samples) for substitution of NAD^+^ with biotin and capture of NAD^+^-capped RNA, and the second aliquot mock-treated (ADPRC-samples) and used as a negative control. Briefly, the aliquots were incubated with 4-pentyn-1-o1 (ThermoFisher Scientific) and ADPRC (Sigma-Aldrich) for ADPRC+ aliquots, or with water for ADPRC-aliquots, 30 minutes at 37 °C. The aliquots were then subjected to biotinylation by copper-catalyzed azide–alkyne cycloaddition by incubation of the RNA with a copper mixture (CuSO_4_, THPTA, Sodium ascorbate, all from Sigma-Aldrich) and biotin azide (Sigma-Aldrich) for 30 minutes at 25 °C. The biotinylated transcripts were captured in Streptavidin Sepharose High Performance (Cytiva). An adenylated adaptor was ligated to the 3’-end of the captured transcripts (Supplementary Table 3) by using a mixture of T4 RNA ligase and truncated T4 RNA ligase 2 (ThermoFisher Scientific and NEB), which served as a template for the reverse transcription primer (Supplementary Table 3). Excess primers were removed by treatment with ExoI (ThermoFisher Scientific), and the cDNA was released by alkaline digest and precipitation. The cDNA was C-tailed for ligation of the second adaptor (named “cDNA anchor” in the original protocol) with Terminal deoxynucleotidyl transferase (ThermoFisher Scientific) and T4 DNA Ligase (ThermoFisher Scientific). Finally, the cDNA was amplified by PCR using Q5 polymerase (NEB) and primers with barcodes for differentiation of the different aliquots and samples (Supplementary Table 3). The products were purified with the AMPure XP kit using a 1X beads:PCR mix ratio (Beckman Coulter). Pooled barcoded libraries (exponential or stationary phase, treated or mock-treated, biological triplicates) were sequenced in a single lane at Illumina HiSeq 2000 in 50 bp single end regime at EMBL Genomics Core Facility, Heidelberg, Germany. Sequencing reads were deduplicated using the clumpify.sh script from the BBMap package v38.46 (https://sourceforge.net/projects/bbmap/), and sequencing adapters were removed using Trimmomatic 0.39 (28). The leading 5’-end G-stretch of variable length was collapsed to a single G using UrQt 1.0.18 (29). That last remaining 5’-end G was then removed, together with 3’-end 7 nt (UMI barcode sequence), and reads shorter than 20 nt were discarded using Cutadapt 1.9.1 (30). Cleaned reads were then aligned to the *M. smegmatis* str. mc^2^ 155 reference genome (NCBI_Nucleotide:NC_008596.1) using HISAT 2.1.0 (31) and SAMtools 1.3.1 (32). Reads with mapping quality ≥10 were kept and inspected using the IGV browser 2.0.23 (33). The reference genome was then divided into open reading frames (ORFs) and intergenic regions (based on NCBI_Assembly:GCF_000015005.1), and reads mapping to each genomic interval were counted using the summarizeOverlaps() function in R/Bioconductor (34). Region read counts were then normalized to library size, pooled across biological replicates, and the ratios of ‘treated’ (ADPRC+) versus the corresponding ‘mock treated’ (ADPRC-) reads were calculated for the areas overlapping the annotated 5’-end of the transcripts (from +1 to +30 positions of the transcript) (35). The reason for limiting the analysis to the TSS-proximal region was that the RNA isolation and conversion protocol yield short fragments of RNA, so only reads mapping close to TSS should represent genuine NAD+-capped RNA. NAD^+^-RNA candidates were defined as those with ≥20 reads in ≥2 biological replicates and a ‘treated’ vs. ‘mock treated’ ratio of ≥3 fold. The complete bioinformatic workflow is available at https://github.com/mprevorovsky/krasny-NADcaptureSeq. The raw sequence read files were deposited in the ArrayExpress database (accession number E-MTAB-10246).

### Sequence analysis

To find common features within the promoter region of the NAD^+^-capped transcripts found in mycobacteria, we retrieved the −50 to −1 sequences of the transcripts detected by NAD captureSeq (IGR-2, IGR-5 and IGR-11 were not included due to uncertainty about their TSS). We then analysed them with the MEME software (36), using multiple motif width settings (default, 4-8, 9-19, and 20-40) and zoops and oops motif site occurrences modes, and the *M. smegmatis* (NC_008596.1) genome sequence as first order Markov model background. We found the GGGTA motif element present multiples times in the MEME results with different parameters; as a representative motif, we chose the motif with the lowest E-value (5.1 e-004) (Figure 5A). Then we used the identified motif to search the sequences, including the IGR-2, IGR-5 and IGR-11 with FIMO with the same background model. Only motif sites with p-val<0.001 were kept. Finally, we used the Jalview software (37) to visualize the sequences with the motif sites.

### Northern blot analysis

RNA for Northern blotting was isolated as described above. Total RNA samples were mixed with stop solution 1:1 (95% formamide, 20 mM EDTA, pH 8.0, 25 mg/ml bromophenol blue, 25 mg/ml of xylene cyanol) and denatured 5 minutes at 95 °C and 1 minute on ice. Aliquots containing 5 μg of RNA were loaded onto 7% PAGE and run at 180 V for 2 h. RNA was transferred to Zeta-Probe blotting membrane (Bio-Rad) in the Trans-Blot SD semi-dry transfer cell (Bio-Rad) at 15 V for 1 h. Membranes were airdried and nucleic acids were then cross-linked by UV exposure on a UVT-20 M transilluminator for 3 minutes and 15 seconds of UV exposure set at high intensity (Herolab). Northern blot pre-hybridization was carried using 6 ml of pre-heated Pre-hybridization Buffer (SET 4X (NaCl 600 mM, Tris-HCl 120 mM, EDTA 8 mM), 0.1 % NaPP, 0.2 % SDS and 500 U/L of heparin solution, Léčiva) for 2 hours at 50 °C. For hybridization, buffer was changed to Hybridization Buffer (SET 4X, 0.1% NaPP, 0.2% SDS, 2,500 U/L of heparin solution and 100 g/L of dextran sulfate), and 5’ ^32^P radiolabelled primers (preparation see below) were denatured by incubating 5 minutes at 95 °C and 1 minute on ice before being added. Hybridization was carried overnight at 50 °C in Techne Hybridizer HB-1D. Finally, membranes were washed sequentially with 6 ml of 2X SSC at room temperature for 15 minutes, 6 ml of 0.1X SSC and 0.1% SDS for 1h at 50 °C, and 6 ml of 0.1x SSC for a few minutes. The membranes were then briefly air-dried, wrapped in plastic foil and set in cassettes for overnight exposure with Fuji MS phosphor storage screens. The image was scanned using an Amersham Typhoon scanner (Cytiva). The images were analysed with QuantityOne software (Biorad), and prepared for presentation using Inkscape free software (https://inkscape.org/).

The oligonucleotide probes were designed using Primer3 online application (https://primer3.org/), and purchased from Eastport or Sigma. 10 pmol of each probe was radiolabelled by incubation with 3 μl of y^32^P ATP (4000 Ci/mmol, 10mCi/ml) (Hartmann Analytic), T4 PNK Buffer 1X and 1 U/μl of PNK (NEB) in a 10 μl final reaction volume at 37 °C for 30 minutes. The probes were then purified using Qiaquick Nucleotide Removal Kit (Qiagen), and eluted in 150 μl of Elution Buffer. Half of this volume was used for each hybridization.

### Ms1 Pull-down assay

To study the NAD^+^ capping of natural Ms1 from *M. smegmatis* cells, we performed Ms1 Pull-down from total RNA. For this, 10 μl of DynaBeads MyOne C1 (ThermoFischer) were sequentially washed with 1 ml of Washing Buffer (5 mM Tris-HCl pH 7.5, 0.5 mM EDTA, 1 M NaCl), two washes with 10 μl of Washing Buffer, two washes with 10 μl of Solution A (0.1 M NaOH, 0.05 M NaCl) and one wash with Solution B (0.1 M NaCl). To attach the biotinylated probe to the beads, 10 μl of 5 μM biotinylated probe (Supplementary Table 3) were added and the beads were incubated for 15 minutes at 25 °C with gentle rotation. The excess probe was removed by washing three times with 10 μl of Washing Buffer. The beads were then resuspended in an equal volume of B&W Buffer (10 mM Tris-HCl pH 7.5, 1 mM EDTA, 2 M NaCl). For Ms1 hybridization to the probe, 10 μl of 1 mg/ml total RNA from *M. smegmatis* exponential or stationary phase cells were added, and the beads were incubated 5 minutes at 65 °C, and then cooled down to 25 °C within 10 minutes. The unbound RNA was removed by washing three times with Washing Buffer, and bound RNA was eluted by incubating the beads in 20 μl of water at 65 °C for 10 minutes twice. Pulled-down RNA was ethanol precipitated and resuspended in water.

### LC–MS data collection and analysis

To screen for the presence of NAD^+^ and NADH, LC–MS was performed using a Waters Acquity UPLC SYNAPT G2 instrument with an Acquity UPLC BEH Amide column (1.7 μm, 2.1mm× 150mm, Waters). The mobile phase A consisted of 10 mM ammonium acetate, pH 9, and the mobile phase B of 100 % acetonitrile. The flow rate was kept at 0.25mL/minutes and the mobile phase composition gradient was as follows: 80 % B for 2 minutes; linear decrease to 68.7 % B over 13 minutes; linear decrease to 5 % B over 3 minutes; maintaining 5 % B for 2 minutes; returning linearly to 80 % B over 2 minutes. For the analysis, electrospray ionization (ESI) was used with a capillary voltage of 1.80 kV, a sampling cone voltage of 20.0 V, and an extraction cone voltage of 4.0 V. The source temperature was 120 °C and the desolvation temperature 550 °C, the cone gas flow rate was 50 L/h and the desolvation gas flow rate 250 L/h. The detector was operated in positive ion mode. For each sample, 8 μL of the dissolved material was injected. Triplicates of Nuclease P1-digested RNA samples were used to identify NAD^+^. Ions with <50 counts were not considered for further analysis. MassLynx software was used for data analysis.

### MRM (multiple reaction monitoring) analysis

The fragmentation studies were performed using SCIEX QTRAP 6500^+^ instrument with an Acquity UPLC BEH Amide column (1.7 μm, 2.1mm× 150 mm, Waters). Mobile phase A was 10 mM ammonium acetate pH 9, and mobile phase B was 100 % acetonitrile. The flow rate was a constant 0.2 mL/minutes and the mobile phase composition was as follows: 80 % B for 2 minutes; linear decrease over 12 minutes to 50 % B; and maintain at 50 % B for 1 minute before returning linearly to 80 % B over 2 minutes. ESI was used with curtain gas of 20 (arbitrary units), Ionspray voltage of 5.5 kV. The ion source gas was 7 (arbitrary units), and the drying gas temperature was 400 °C. The declustering potential was ^+^80 V, the entrance potential ^+^10 V, collision energy ^+^40 V. The detector was operated in positive ion mode. For the quantification of NAD^+^, the first precursor ion was [M ^+^ H]^+^ at m/z 664.108 and the second ions selected were [M ^+^ H]^+^ at m/z 428.00 and 524.00. For each sample, 8 μL of the dissolved material was injected.

### Mass spectrometry result quantification

A total of 2 mg of total RNA from each organism was divided into four aliquots. After digestion with Nuclease P1 (NuP1), each aliquot was spiked with increasing concentration of NAD^+^ (0 μM, 15 μM, 50 μM, 150 μM). After MRM (multiple reaction monitoring) analysis of all the aliquots, the area under the peak was calculated and plotted against the concentration. The experimental points were fitted with a linear regression and the intercept with X-axis represents the concentration of the cap in the sample. The quantification was repeated three times and the final value was obtained from the average of the three measurements with relative SD.

### Calculation of NAD^+^ cap amount in Ms1

For the estimation of the amount of NAD^+^ capped Ms1 15.6 ng (155 fmol) of purified Ms1 from *M. smegmatis* from stationary phase were digested with NuP1 and analysed with SCIEX QTRAP 6500^+^. Using a calibration line 37.9 fmol of NAD^+^ were detected in the samples that correspond to 24.5 % of RNA having NAD^+^ caps. Since the Ms1 was 85 % pure, the final amount of the capped Ms1 was 28.2 %.

### Production of control transcripts

RNAI and A-less m^7^G-capped transcripts were used as controls for some experimental procedures. The transcription template for RNAI was produced by using p770 plasmid (from *E. coli* strain LK_2385) as template for PCR amplification using Q5 polymerase (NEB) and LK_3413 and LK_3414 primers to introduce a T7 class II promoter (Supplementary table 3). A-less template for transcription was produced by using LK_2756 and LK_2757 on LK_2753 as PCR template. The product was purified and used as a template for amplification with LK_2797 and LK_2798 to introduce EcoRI and HindIII restriction sites in order to clone A-less control transcript into p770 plasmid to transform *E. coli* with (strain LK_2346). The plasmid was isolated and used as a template for PCR amplification with LK_3331 and LK_3332 to introduce a T7 class II promoter. All plasmids were isolated using QIAprep Spin Miniprep Kit (Qiagen), and all PCR products were purified with QIAquick Gel Extraction Kit (Qiagen). RNAI and A-less transcription templates with T7 class II promoters were used for *in vitro* transcription with the Promega Ribomax Transcription System as specified by the manufacturer. The product was m^7^G-capped using the Vaccinia Capping System (NEB). To eliminate any uncapped remnant, we treated these samples with 5’ polyphosphatase (Lucigen) (20 U per 5 μg of RNA, 1h of incubation at 37 °C), and T erminator 5’-Phosphate Dependent Exonuclease (Lucigen) (1 U per 5 μg of RNA, 1h of incubation at 30 °C using Buffer A). Before each treatment RNA was denatured by incubating at 65 °C 5 minutes and on ice for 5 minutes before each treatment. After each treatment, RNA was purified by sequential phenol:chloroform extractions and ethanol precipitation.

### Ms1 capture and reverse transcription

Samples containing 12.5 μg of RNA were divided into 2 aliquots that were treated or mock treated with ADPRC. Briefly, RNA aliquots in 10 μl volumes were mixed with 20 μl of 125 μg/ml of ADPRC (ADPRC+, enzymatic treatment) or H_2_O (ADPRC-, mock treatment), 20 μl of ADPRC Reaction Buffer 5x (250 mM Na-HEPES pH 7.0 and 25 mM MgCl_2_), and 10 μl of 100 % pentynol in a final reaction volume of 100 μl, incubated for 30 minutes at 37 °C, extracted sequentially with equal volumes phenol / phenol-choroform 1:1 / chloroform, and precipitated by adding of 2 μl of glycogen, 0.1 volumes of 3 M sodium acetate, and 2.5 volumes of 100 % ethanol, incubated for 1 hour at −80 °C and centrifuged for 1 hour at 4 °C and 32,700 g. The pellet was then washed twice with 75 % ethanol and air-dried for 5 minutes before being resuspended in 85 μl of 1X ADPRC Reaction Buffer. The RNA was then mixed with 12.5 μl of 2 mM biotin azide and 3.4 μl of copper mix (CuSO_4_ 1 mM, THPTA 0.5 mM, and Sodium Ascorbate 2 mM), and incubated 30 minutes at 25 °C while shaken at 350 r.p.m. RNA was then again extracted and precipitated as described above, and the RNA pellets were resuspended in 41 μl of Immobilization Buffer (10 mM Na-HEPES pH 7.2, 1 M NaCl and 5 mM EDTA). 50 μl of resuspended streptavidin sepharose (Cytiva, cat. no. 17-5113-01) were added into columns (Mobitec, cat. no. M1003, with 10 μm filters from Mobitec), and equilibrated by washing 3 times with 200 μl of Immobilization Buffer and centrifuging 1 minute at 16,000 g at room temperature. The beads were blocked by incubation with 100 μl of Immobilization Buffer supplemented with 100 μg/ml of acetylated BSA for 20 minutes at 20 °C and shaken at 800 r.p.m., and washed 3 times with 200 μl of Immobilization Buffer. Then the beads were incubated with the RNA aliquots for 1 hour at 20 °C and shaken at 800 r.p.m., and washed 5 times with 200 μl of Streptavidin Wash Buffer (50 mM Tris-HCl pH 7.4, and 8 M urea). The beads were subsequently equilibrated by washing 3 times with 200 μl of First-strand Buffer 1X (50 mM Tris-HCl pH 8.3, 75 mM KCl and 3 mM MgCl_2_), and blocked by incubation with 100 μl of First-strand Buffer supplemented with 100 μg/ml of acetylated BSA for 20 minutes at 20 °C and shaken at 800 r.p.m. The beads were finally washed 3 times with 200 μl of First-strand Buffer 1X. Reverse transcription of captured transcripts was carried by addition of 1.5 μl of 10 μM random hexamers as non-specific primers (Eastport), 1.5 μl of 10 mM dNTP mix, and 18.5 μl of H_2_O, denaturation at 65 °C for 5 minutes and incubating on ice 5 minutes, and subsequent addition of 1.5 μl of 100 mM DTT, 6 μl of First-strand Buffer 5X, and 1 μl of Superscript III (Invitrogen). The columns were incubated 5 minutes at 25 °C, 1 hour at 50 °C, and 20 μl of H_2_O were added before the final enzyme inactivation of 15 minutes at 70 °C. The columns were centrifuged, and cDNA was eluted twice with 100 μl of H_2_O. A final incubation with 100 μl of 150 mM NaOH was performed for 25 minutes at 55 °C to ensure the release of all cDNA, and the pooled eluate was ethanol precipitated as described above. The RNA was resuspended in H_2_O and used for RT-qPCR quantification of Ms1 and the m^7^G-capped negative control.

### Enzymatic treatment

Capped transcripts have increased resistance to 5’-phosphatases compared to their uncapped counterparts (7, 38, 39). We confirmed that NAD^+^-capped transcripts were approximately as resistant as their m^7^G-capped form, and that both of them were at least ~10 times more resistant than their uncapped form (data not shown). To measure the resistance of Ms1 to the enzymatic treatment in different growth phases, we spiked biological triplicates of *M. smegmatis* cells at exponential and stationary phases of growth with an m^7^G-capped control (Figure 3A). After isolation of total RNA, samples of 12.5 μg of RNA were divided into two aliquots. Each aliquot was either enzymatically treated with 5’ polyphosphatase (Lucigen) (20 U per 5 μg of RNA, 1h of incubation at 37 °C) and Terminator 5’-Phosphate Dependent Exonuclease (Lucigen) (1 U per 5 μg of RNA, 1 h of incubation at 30 °C using Buffer A), or mock treated (treated only with Terminator 5’-Phosphate Dependent Exonuclease) (Figure 3B). Before each treatment RNA was denatured by incubating at 65 °C for 5 minutes and on ice for 5 minutes. After each treatment, RNA was purified using the Clean-up and Concentration kit (Norgen). The RNA left in each aliquot was reverse transcribed for 1 h with Superscript III (Invitrogen) and random hexamers as non-specific primers (Eastport), and Ms1 and the m^7^G-capped control were quantified by RT-qPCR with SYBR Green (Sigma). The fraction of treatment-resistant Ms1 was calculated by dividing the amount of transcripts left after the enzymatic treatment (resistant) by the amount left after the mock treatment (total), and normalizing to the m^7^G-capped control. The result shows that Ms1 from *M. smegmatis* cells at stationary phase is more resistant to the enzymatic treatment than Ms1 from cells at exponential phase (Figure 3C).

### Purification of *Mycobacterium smegmatis* RNAP core

*Escherichia coli* DE3 expression strain containing a plasmid pRMS4 coding for subunits of the RNAP core (LK_1853; from Kouba *et al.,* 2019) was grown at 37 °C to exponential phase (OD_600_ ~ 0.5). Expression of RNAP was induced with 500 μM IPTG for 4 hours at room temperature. Cells were harvested by centrifugation, washed, resuspended in P Buffer (300 mM NaCl, 50 mM Na_2_HPO_4_, 5 % glycerol, 3 mM 2-mercaptoethanol) and disrupted by sonication. Cell debris was removed by centrifugation and supernatant was mixed with 1 ml Ni-NTA Agarose (Qiagen) and incubated for 90 minutes at 4 °C with gentle shaking. Ni-NTA Agarose with bound RNAP was loaded on a Poly-Prep Chromatography Column (BIO-RAD), washed with P Buffer and afterwards washed with P Buffer supplemented with 30 mM imidazole. The protein was eluted with P Buffer containing 400 mM imidazole and fractions containing RNAP were pooled and dialyzed against Storage Buffer (50 mM Tris-HCl, pH 8.0, 100 mM NaCl, 50 % glycerol, 3 mM 2-mercaptoethanol). The isolated RNAP complex was stored at −20 °C.

#### Purification of *Mycobacterium smegmatis* Sigma A

*Escherichia coli* DE3 expression strain containing a plasmid pET22b with the gene for *M. smegmatis* σ^A^ (LK_1740 (41)) was grown at 37 °C until OD_600_ reached ~ 0.5. Expression of σ^A^ was induced with 300 μM IPTG for 3 hours at room temperature. Isolation of σ^A^ was performed in the same manner as RNAP purification with one difference. Instead of using P Buffer with 30 mM imidazole to wash Ni-NTA Agarose with bound σ^A^, P buffer with 50 mM imidazole was used. The protein was then eluted with P Buffer containing 400 mM imidazole but instead of the purification on a column, batch purification and centrifugation were used to separate the matrix and the eluate. Fractions were then dialyzed and stored in the same manner as it was done for RNAP.

### Abortive transcription

*M. smegmatis* RNAP and σ^A^ were mixed at a 1:20 ratio to a final concentration of the RNAP holoenzyme of 500 nM and incubated at 37 °C for 30 minutes. 2 μl of the RNAP holoenzyme were then mixed with 250 ng of plasmid containing the sequence of Ms1 under its native promoter (LK_2166), 0.5 μl of KCl 1M, 0.1 μl of 100X BSA, 40 mM Tris-HCl pH 8.0, 10 mM MgCl_2_, 1 mM dithiothreitol (DTT), and water to a final volume of 8 μl. This mixture was incubated at 37 °C for 10 minutes to allow open complex formation. Then, to allow the transcription reaction to proceed until position ^+^2 (Figure 4), we added 2 ul of the nucleotide mix containing the following components (final concentrations in the 10 μl reaction mix are indicated in parentheses); CTP (50 μM), α-^32^P-CTP from Hartman analytics, 10mCi/ml (100-fold diluted), ATP (200 μM when present), NAD^+^ (200 μM when present). The tubes were incubated at 37 °C for 10 minutes and the reaction was stopped by the addition of 10 μl of formamide stop solution supplemented with amaranth red, and 5 μl of the reaction was run in 20 % polyacrylamide sequencing gel at 1,800 V for 1.5-2h. The gels were dried and exposed to Fuji MS phosphor storage screens and scanned with an Amershan Typhoon scanner (Cytiva). The signals were quantified using Quantity One software (Bio-Rad), and normalized to the last sample (set as 1), which only contains NAD^+^ with no ATP.

## RESULTS

### NAD^+^-RNA detection in Archaea and mycobacteria

To determine whether chemical modifications are present at the 5’-end of RNA in Archaea and mycobacteria, we performed mass spectrometry analysis of total RNA samples from the methanogenic archaeon *M. barkeri* and the mycobacterial *M. smegmatis* as previously described (2, 38). The MRM analysis revealed the presence of NAD^+^ covalently attached to RNA in both organisms (Figure 1, Supplementary Figure 1A). Namely, the appearance of two characteristic peaks on the chromatogram, corresponding to fragments of NAD^+^, in RNA samples digested with nuclease (+NuP1, Figure 1A, B, C and D) confirmed the presence of NAD^+^, whereas the absence of these peaks in the negative control samples (i.e. sample not treated with nuclease, -NuP1) indicated that NAD^+^ was covalently attached to the RNA. We found that the NAD^+^ level in *M. barkeri* in stationary phase of growth was 7.4±0.5 fmol per μg of total RNA (Figure 1A and E). Likewise, the presence of NAD^+^ was assessed in total RNA isolated from *M. smegmatis* cells from exponential and stationary phases of growth. While in exponential phase the detected NAD^+^ capping level was relatively low (2.4±0.5 fmol/μg of total RNA), it increased more than 50-fold in stationary phase (116±37.5 fmol/μg of total RNA) (Figure 1B, C and E). In addition, we also detected NADH, which was covalently bound to RNA in both organisms, albeit at much lower concentrations: approximately 0.006 fmol/μg of total RNA in *M. barkeri* and 22.5 fmol/μg of RNA in *M. smegmatis* in stationary phase (Figure 1E, Supplementary Figure 1B). As the LC-MS conditions were optimized mainly for NAD^+^, other RNA modifications, namely dephospho-Coenzyme A (dpCoA) and Flavin adenine dinucleotide (FAD), were not detected (42–45). The incidence of NAD^+^ capping in RNA from Archaea and exponentially growing mycobacteria is comparable to the levels detected in bacteria, dengue virus, plant and mammalian cells (2, 4, 8, 14, 45), and the higher level of NAD^+^ capping in *M. smegmatis* in stationary phase are comparable to those reported for *E. coli* and S. *cerevisiae* (46) (Figure 1F). These findings show that NAD^+^ capping exists in all domains of life (Figure 1G).

**Figure 1.**
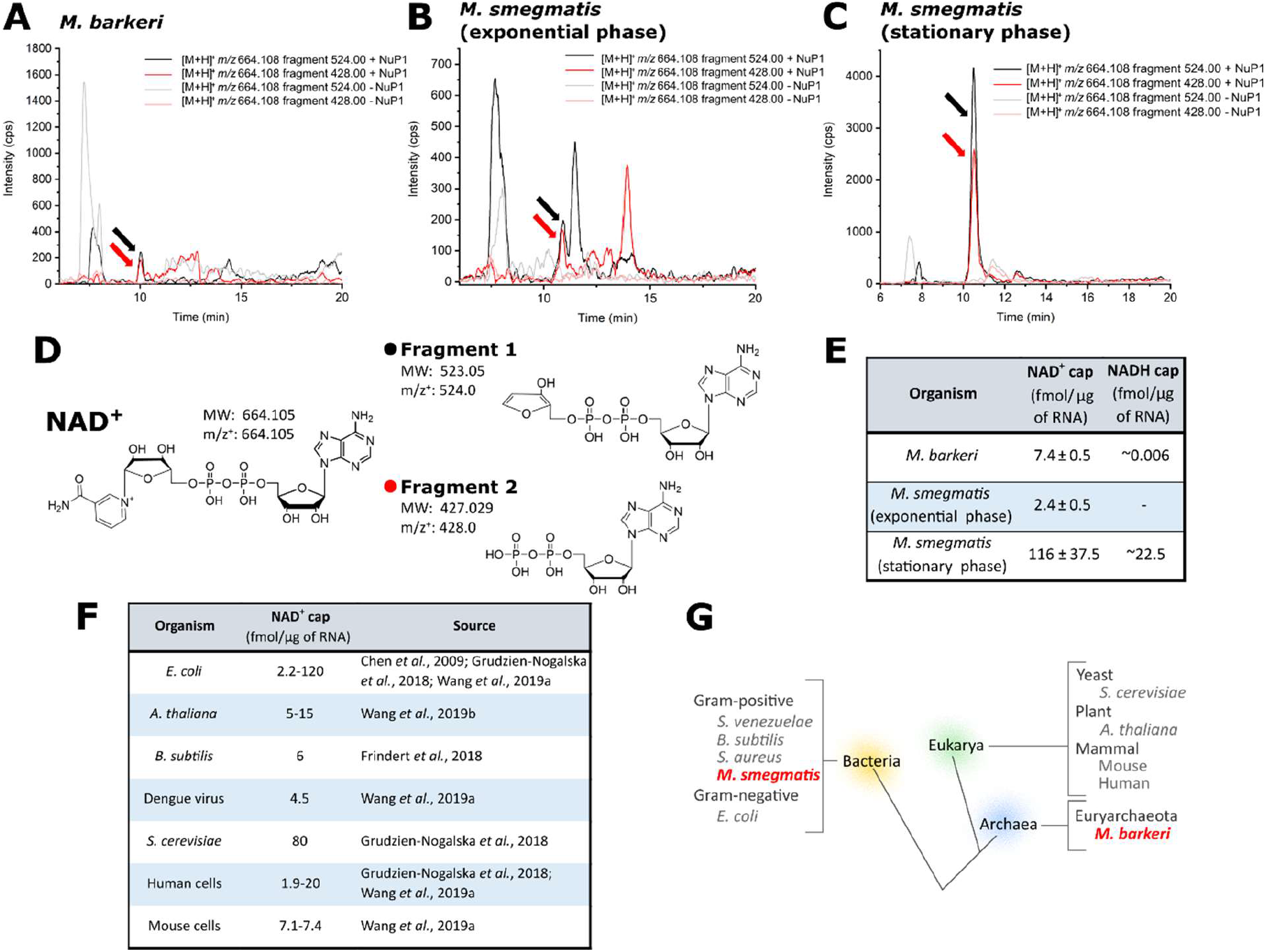
NAD^+^ caps exist in all domains of life; detection of NAD^+^ and NADH caps in Archaea and mycobacteria. A.B.C. Examples of MRM Chromatograms showing detection of two fragments of NAD^+^ in total RNA digested with Nuclease P1 (+NuP1, black and red lines) compared to the negative control (-NuP1, grey and pink lines) in samples extracted from *M. barkeri* (**A**), and *M. smegmatis* in exponential phase (**B**) and stationary phase (**C**). **D**. Structure, molecular weight (MW) and mass/charge (m/z) of NAD^+^ (left) and the two fragments detected by MRM analysis (right), indicated with black arrows (m/z 524) or red arrows (m/z 428) in the chromatograms (**A**, **B** and **C**). **E**. Quantified levels of NAD^+^ (measured in triplicate) and approximate levels of NADH covalently attached to total RNA of *M. barkeri,* and *M. smegmatis* in exponential and stationary phases of growth. **F**. Approximate values of previously reported levels of NAD^+^ capping in bacteria and eukaryotic cells (2, 4, 45, 46). **G**. Schematic tree of life showing all organisms where NAD^+^ capping has been detected; the discoveries presented in this work are highlighted in red.

### Identification of NAD^+^-capped transcripts in *M. smegmatis*

To further characterize NAD^+^ capping and identify the NAD^+^-capped transcripts in mycobacteria, we applied NAD captureSeq (27, 47) to biological triplicates of total RNA samples from *M. smegmatis* cells harvested from exponential and stationary growth phases. Briefly, nicotinamide residues of the NAD’-RNA within total RNA were substituted with biotin (a process hereafter referred to as biotinylation). The biotinylated RNAs were selectively captured via biotin-streptavidin interaction. The captured transcripte were sequenced and their reads were compared to those obtained from non-biotinylated negative control samples (27, 47). We detected 25 transcripts with ≥20 reads in ≥2 biological replicates, ≥3 fold enrichment over non-biotinylated negative control that overlapped with reported TSS of the transcript (23, 35)). Of the 25 transcripts, 21 contained open reading frames (ORFs) (Supplementary Table 1). The gene products encoded by the captured mRNAs are involved in cell envelope functions and redox processes (Table 1). Interestingly, the captured transcripts included the well-characterised small RNA. (sRNA) Ms1 (23, 48). In addition, three other validated but functionally uncharacterized *M. smegmatis* sRNAs were detected by NAD captureSeq (49, 50) (Table 2).

**Table 1.** Function of NAD^+^-RNAs in *M. smegmatis.* The most frequently repeated ontology terms of the transcripts detected by NAD captureSeq in *M. smegmatis.* The ontology terms of the NAD^+^-RNAs were collected using QuickGO database from EMBI-EBI (www.ebi.ac.uk/QuickGO/).

| GO:0016020 | GO:0016021 | GO:0016491 | GO:0005737 | GO:0046872 |
| --- | --- | --- | --- | --- |
| Membrane | Integral component of membrane | Oxidoreductase activity | Cytoplasm | Metal ion binding |
| MSMEG_6345 | MSMEG_6345 | MSMEG_0670 | MSMEG_1919 | MSMEG_1919 |
| MSMEG_6676 | MSMEG_6676 | MSMEG_5246 | MSMEG_1605 | MSMEG_1112 |
| MSMEG_0987 | MSMEG_0987 | MSMEG_4268 | MSMEG_1519 | MSMEG_4268 |
| MSMEG_3689 | MSMEG_3689 |  |  |  |
| MSMEG_5648 | MSMEG_5648 |  |  |  |
| MSMEG_4268 | MSMEG_4268 |  |  |  |

**Table 2.** NAD^+^-sRNAs in *M. smegmatis*. The fold enrichment at the 5’-end of sRNAs detected by the NAD captureSeq experiment (compared to non-biotinylated negative controls) in *M. smegmatis* in exponential (Exp.) and stationary (Stat.) phases, respectively. The orientation of the sRNAs and their adjacent genes are indicated with arrows, the sRNA is in bold.

| Name | Adjacent genes and their orientation | NAD captureSeq enrichment |  | Source | Rfam term |
| --- | --- | --- | --- | --- | --- |
|  |  | Exp. | Stat. |  |  |
| Ms1 | MSMEG_6173 -<br>MSMEG_6174<br>→ → ← | - | 4.0 | Hnilicova et al., 2014;<br>Šíková et al., 2019 | RF02566 |
| IGR-2 | MSMEG_0373 -<br>MSMEG_0374<br>→ → ← | - | 7.6 | Li et al., 2013 | RF02467 |
| IGR-5 | MSMEG_5794 -<br>MSMEG_5795<br>→ ← ← | 5.9 | Inf. | Li et al., 2013 | RF02471 |
| IGR-11 | MSMEG_0358 -<br>MSMEG_0359<br>→ ← ← | 3.9 | 31.1 | Li et al., 2013 |  |

Further analysis of the +1 position of all detected transcripts revealed that 18 out of the 21 transcripts containing an ORF had a +1A, a requirement for NAD^+^ capping (15, 35). Although the other three transcripts did not contain a +1A, we noticed that the sequencing signal perfectly matched positions that encoded an A only two nucleotides away from the annotated TSS (Supplementary Table 1, Supplementary Figure 2A). This observation suggests that these transcripts may have alternative TSS, as it is often the case in *M. smegmatis* (51).

Next we compared the reported TSSs of Ms1 and the three uncharacterized sRNAs (IGR-2, IGR-5 and IGR-11) detected by NAD captureSeq with the TSSs inferred from our sequencing data. In the case of Ms1, the reported +1A TSS perfectly matched the NAD captureSeq signal (position 6242368, Figure 2A) (23, 48). In contrast, the previously reported TSSs of IGR-5 and IGR-11 (transcribed from the intergenic regions between MSMEG_5794 - MSMEG_5795, and MSMEG_0358 - MSMEG_0359, with reported TSSs at positions 5862669 and 396650, respectively (Figure 2C and D) (49)) did not contain A. These previously reported TSSs are 30 and 181 nucleotides away from the TSSs detected in our study. Importantly, the TSSs for IGR-5 and IGR-11 detected in our study contain A (positions 5862639 and 396831, respectively). This is consistent with a previous RNA-Seq experiment that showed transcriptional signals downstream of these positions (48). As for IGR-2, which is transcribed from the intergenic region between MSMEG_0373 and MSMEG_0374 (Figure 2B) (49), the reported TSS in position 417847 does not encode an A. However, our sequencing signal begins 32 nucleotides away (at position 417879) with an A.

**Figure 2.**
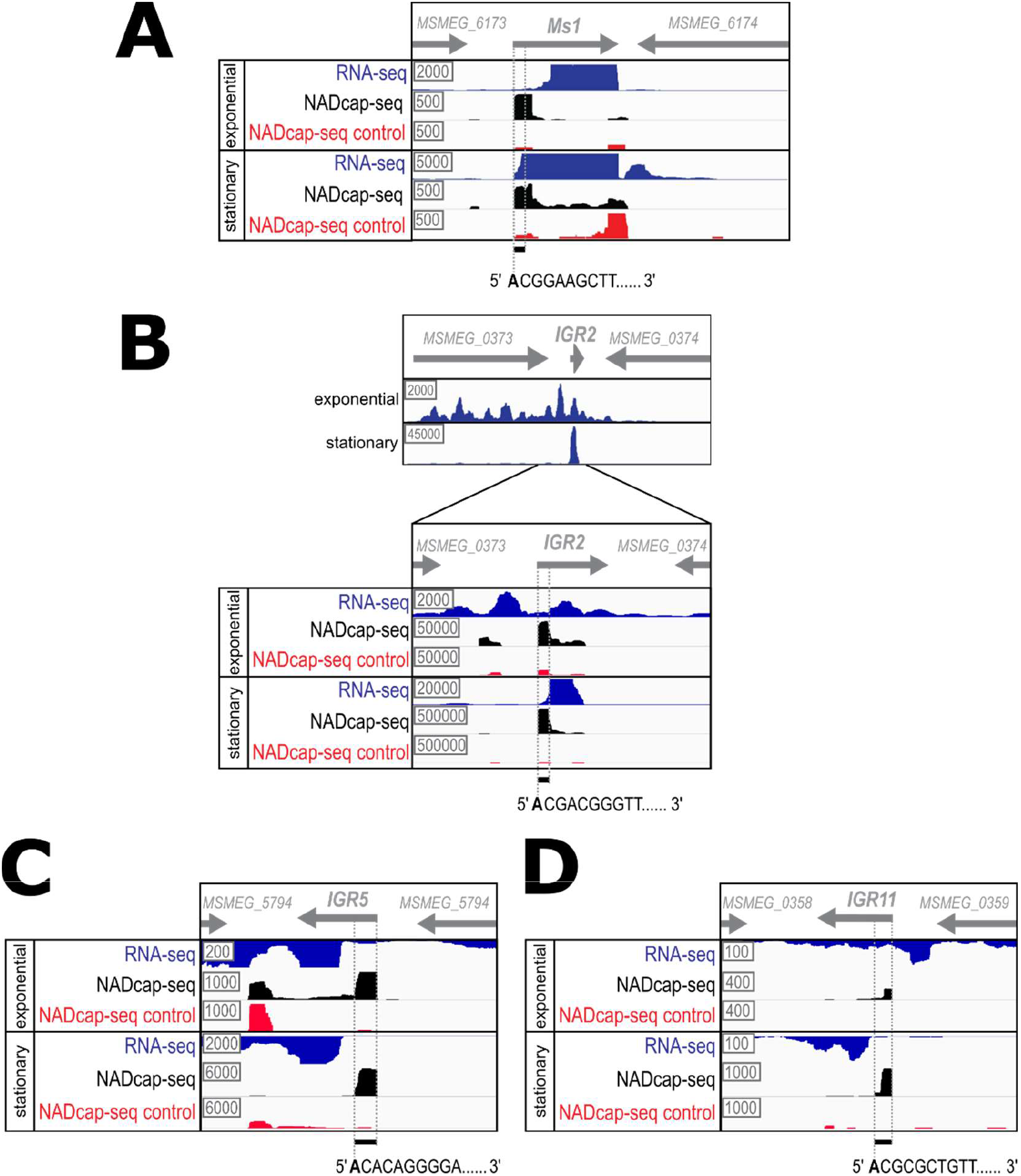
NAD captureSeq detection of sRNAs in *M. smegmatis*. A combination of previous RNA-seq data (blue) (48), data from the NAD captureSeq experiment (black) and NAD captureSeq negative control (red) from this study are shown, visualized by IGV software. Representative biological replicates from exponential and stationary phases of growth are shown. The grey arrows in each panel indicate the orientation of adjacent genes and detected sRNAs. The +1 position as detected by the NADcaptureSeq experiment and the +1 to +30 regions used for 5’-end enrichment calculation are indicated with dashed lines. The nucleotide sequences of the 5’ ends of the respective NAD captureSeq signals with detected TSSs (in bold) are indicated below. **A**. Ms1; **B.** IGR-2, **C.** IGR-5, **D.** IGR-11.

Considering the differences between the reported TSSs and the beginning of the NAD captureSeq signal, we analysed the transcription pattern of the highly expressed IGR-2 transcript by Northern blotting. The experiments revealed that there are at least two variants of IGR-2 (Supplementary Figure 2A and B) of approximately 83 nt and, the more prominent one, 123 nt, suggesting the presence of alternative TSSs at least for IGR-2.

To analyse whether the detection of the captured transcripts may have been biased by their higher abundance *in vivo,* we analysed the expression level of the 25 transcripts detected by NAD captureSeq in a previous RNA-Seq study of *M. smegmatis* (48). The transcripts captured by NAD captureSeq showed a wide range of expression levels, spanning four orders of magnitude. However, expression of many captured transcripts was higher than the median expression value of the transcripts detected by RNA seq (Supplementary Figure 3A). To find out whether the enrichment of transcripts detected in NAD captureSeq correlated with the expression levels of these transcripts in the cell, we plotted the enrichment of captured transcripts detected by NAD captureSeq as a function of the logarithm of their transcript levels detected by RNA-Seq (Supplementary Figure 3B) (48). The resulting plot revealed no proportionality between the two functions, as compared to an example of perfect direct proportionality between Y and X.

### Ms1 sRNA is NAD^+^-capped

Ms1 is a non-coding sRNA that interacts with the RNAP core in *M. smegmatis,* affecting its intracellular level, and it is the most abundant non-ribosomal RNA of *M. smegmatis* in stationary phase (23, 48). Hence, we selected it to validate the NAD^+^ captureSeq results. To confirm the presence of NAD^+^ cap on Ms1, we first pulled down Ms1 from total RNA of *M. smegmatis* from exponential and stationary growth phases and analysed the samples by mass spectrometry using MRM analysis. While we did not detect any significant signal for NAD^+^ in the Ms1 sample obtained from cells in exponential phase, we detected ~2.4 pmol of NAD^+^ per μg of RNA in Ms1 from stationary phase cells (Supplementary Figure 4), suggesting that approximately 30 % of cellular Ms1 bears an NAD^+^-cap in this phase of growth.

Next, we validated the NAD^+^ capping of Ms1 by combining the biotinylation approach with RT-qPCR quantification (a procedure also known as “NAD-qPCR”), similarly as described in other studies (3, 4, 6, 10, 52). Briefly, we subjected total *M. smegmatis* RNA samples extracted from stationary phase, spiked with an m^7^G-capped transcript as a negative control, to substitution of NAD^+^ with biotin (or mock treatment without substitution), and subsequent capture by streptavidin beads followed by RT-qPCR. This experiment showed that Ms1 was approximately six times more frequently captured in the treated sample compared to the mock treated sample, while the negative control was not (Supplementary Figure 5).

Finally, we validated the capping of Ms1 by a third approach, based on enzymatic treatment. We took advantage of the increased resistance of NAD^+^-capped transcripts to 5’-phosphatase compared to their uncapped counterparts, which renders 5’ monophosphorylated transcripts susceptible to 5’-3’ exonucleolytic degradation, whereas those with 5’ triphosphates or NAD^+^ caps are not (3, 53). We spiked samples of *M. smegmatis* cells from exponential and stationary phases of growth with an m^7^G-capped control, and extracted total RNA (Figure 3A). We then subjected RNA samples to the enzymatic treatment (Figure 3B). Ms1 was quantified by RT-qPCR and normalized to the m^7^G-capped control. The experiment revealed that Ms1 was resistant to the enzymatic, especially when extracted from stationary phase cells (Figure 3C). Taken together, these results strongly indicate that Ms1 is, indeed, NAD^+^-capped, and that its capping level is higher in stationary phase relative to exponential phase.

**Figure 3.**
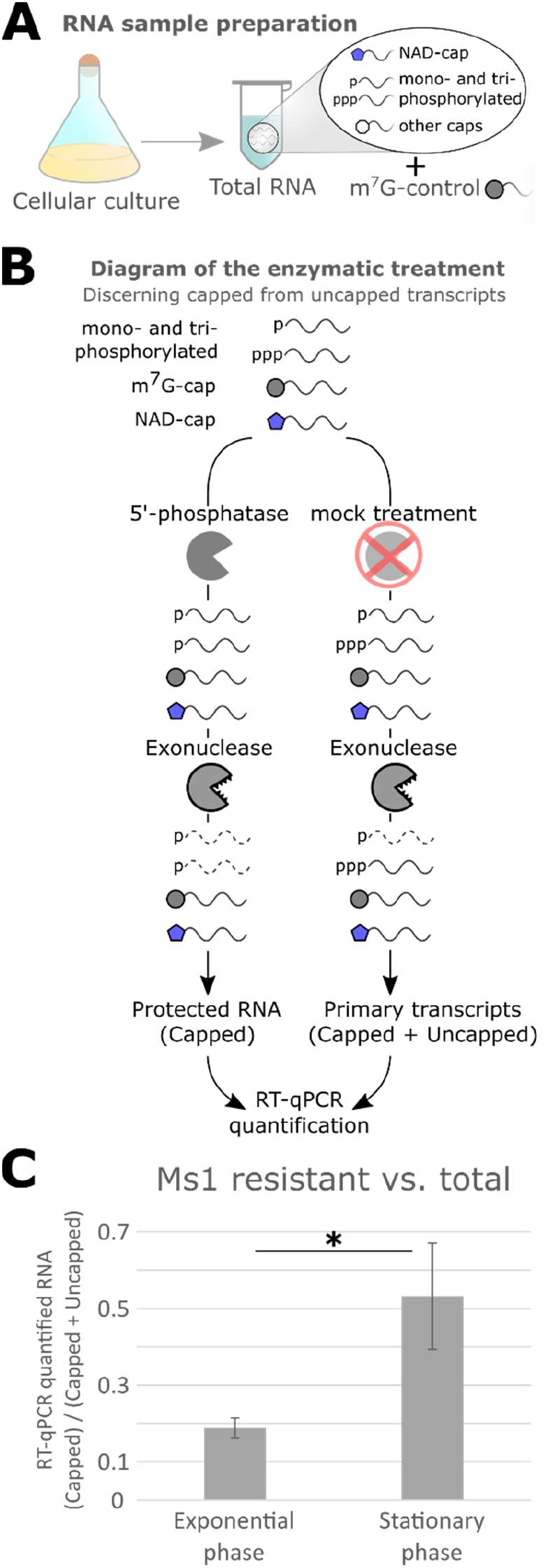
NAD^+^ capping of Ms1 sRNA in exponential and stationary phases of growth. **A**. Total RNA samples from *M. smegmatis* cells from exponential and stationary phases were spiked with a m^7^G-capped control transcript for quantification purposes (15). **B**. Diagram of the enzymatic treatment: 5’-phosphatase turns triphosphorylated RNA into mono-phosphorylated RNA, which can be then degraded by a 5’-3’ exonuclease. Capped transcripts are less sensitive to 5’-phosphatase, and therefore more resistant to the combined enzymatic treatment (53). **C**. The RNA samples were enzymatically treated or mock treated (see panel B and Materials and Methods section), and Ms1 was quantified by RT-qPCR and normalised to the m^7^G-capped control. The plot shows the ratio between Ms1 detected after enzymatic treatment (Capped) and mock treatment (Capped + Uncapped), in RNA from exponential and stationary phases of growth. The bars show average values, the error bars ±SD (p=0.035; t-test type 1, 1 tail, n=3).

### Mycobacterial RNAP can initiate transcription of Ms1 with NAD^+^ *in vitro*

The ability of the RNAP of T7 phage, *E. coli, B. subtilis, S. cerevisiae* and mitochondrial RNAP from human and yeast to initiate transcription with NAD^+^ has been demonstrated in the last few years (2–4, 11, 12, 15, 43, 44, 54), but to date the actinobacterial RNAP has not been tested for this ability. Therefore, we employed the abortive transcription approach as previously described (15, 55), using the natural promoter of Ms1 to initiate transcription of the Ms1 gene (Figure 4) (23, 48). As the Ms1 promoter was previously suggested to initiate transcription with σ^A^ *in vivo* (23), we used *M. smegmatis* RNAP associated with this σ factor. Transcription reactions were carried out in parallel using different defined ratios of NAD^+^ to ATP (Figure 4). The NAD^+^-capped abortive transcripts were quantified (NAD^+^-C*) and plotted. The graph shows that transcription from the Ms1 promoter was efficiently initiated with NAD^+^ (lane 3) and that the equimolar presence of ATP to NAD^+^ reduced NAD^+^ capping by ~60 %.

**Figure 4.**
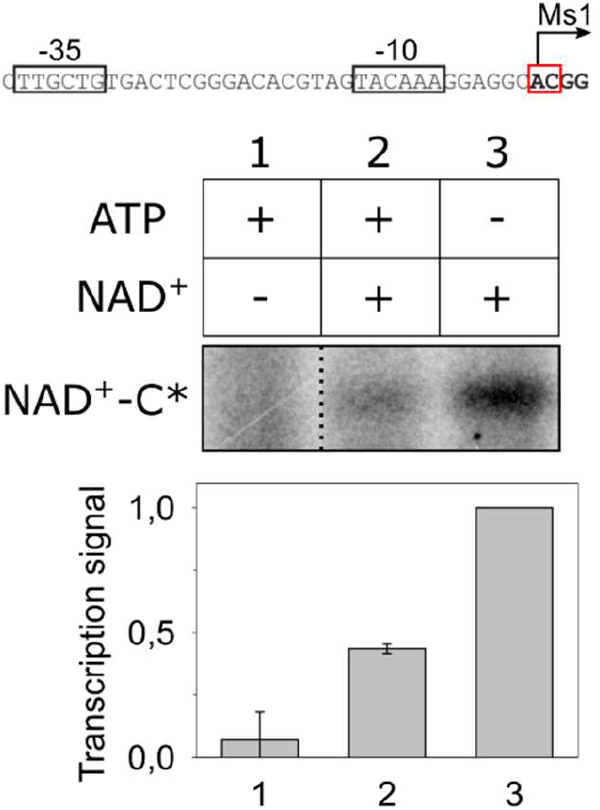
RNAP from *M. smegmatis* incorporates NAD^+^ caps into Ms1 *in vitro.* Abortive transcription was used to monitor the relative affinities of ATP and NAD^+^ as initiating substrates for RNAP at the Ms1 promoter (shown on top). The +1A and +2C positions are shown in bold in the red box. Representative primary data show the abortive transcript (NAD^+^-C*) formed by NAD^+^ and the radiolabelled CTP (C*). The presence/absence of ATP and NAD^+^ (each 200 μM) is indicated. The dashed line indicates that the lanes from the same gel were electronically assembled for presentation. The graph shows the quantification of the NAD^+^-C* dinucleotide, normalized to the transcription signal in the absence of ATP, which was set as 1 (lane 3). The bars show average values ± SD, n=3.

### Promoters of NAD^+^-capped targets are potential members of σ^A^ and σ^F^ regulons

Previous studies found that the promoter sequence determines the NAD^+^ capping incidence (15, 53). To find out whether mycobacterial NAD^+^-RNAs have common promoter sequence motifs, we analysed the promoter regions of all the genes detected by NAD captureSeq in our study (Supplementary Table 1). We found a 27 bp long motif in 18 sequences. The motif contained two conserved regions: (i) GGGTA and (ii) GNT (Figure 5A). This motif is fully or partially present in 19 out of the 25 genes (Figure 5B) and is highly reminiscent of the σ^F^ consensus sequence (Figure 5B). Visual inspection of the promoter regions then revealed three sequences that did not match the σ^F^ consensus sequence (Figure 5B). One of them was Ms1, which is σ^A^ dependent (Figure 5C). This suggests that the NAD^+^ capped transcripts originate either from σ^F^ or σ^A^ dependent promoters.

**Figure 5.**
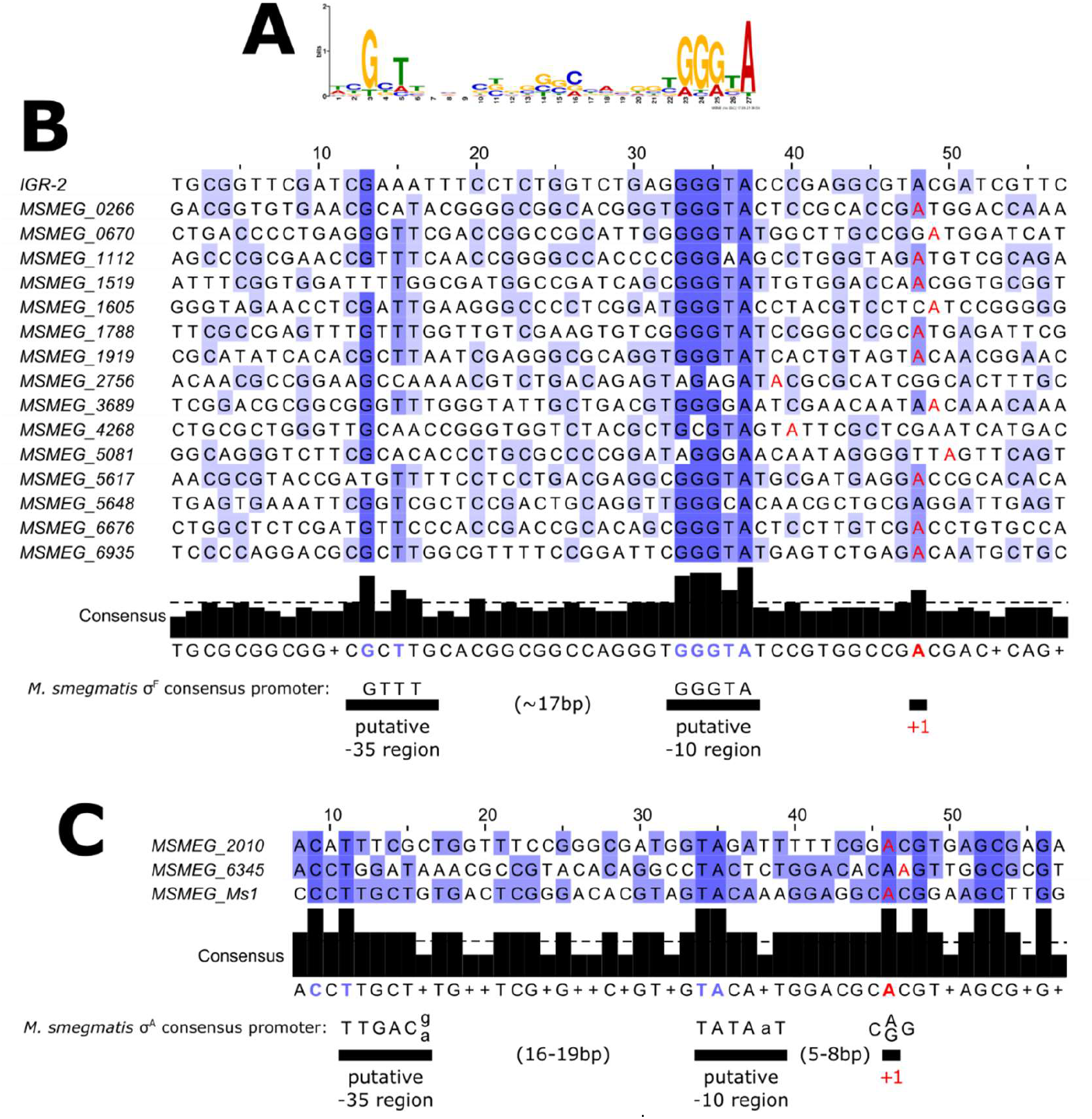
Sequence alignement of promoter regions of NAD^+^-RNAs in mycobacteria. **A**. Sequence logo found in TSS-upstream regions of transcripts detected by NAD captureSeq. **B**. Alignment of promoter sequences containing the consensus sequence for σ^F^. The alignment is centred at the putative −35 region (conserved GNT in the logo). The graph below indicates the level of conservation, the dashed line indicates 50% conservation. The >50 % conserved positions are showed in blue, the +1A positions detected by our study are showed in red. and the consensus −35 and −10 regions of *M. smegmatis* σ^F^ dependent promoters are shown for comparison (51). **C.** Alignment of promoter sequences containing the consensus sequence for σ^A^ recognition. The graph below shows the conservation; *M. smegmatis* σ^A^ consensus sequence is shown below the graph (56).

## DISCUSSION

NAD^+^ capping is an RNA modification that occurs in prokaryotic and eukaryotic cells and for which the biological roles have not been clearly defined. Its nature suggests that it may be a universal modification present in all living organisms. In this work we searched for NAD^+^ caps in the Archaeal kingdom of life, as well as in the still unexplored clinically relevant Mycobacterium genus. The analysis of RNA from archaeal *(M. barkeri)* and mycobacterial *(M. smegmatis)* cells by mass spectrometry has revealed that NAD^+^ is covalently attached to RNA in both organisms, confirming the existence of NAD^+^ caps in all domains of life. This result, along with previous findings, suggests the evolutionary conservation of NAD^+^capping in all three domains of life. Moreover, the incidence of NAD^+^ capping in archaeal and mycobacterial RNA is similar to that of mammalian cells, bacteria and dengue virus (2, 4, 45, 46). The observed variation in NAD^+^ capping levels in *M. smegmatis* (50-fold increase in stationary phase cells) resembles previous findings from *E. coli,* where the NAD^+^ capping level of RNAI sRNA doubled in stationary phase (15), correlating with a decrease in intracellular ATP concentration and an increase in intracellular NAD^+^ concentration in this phase of growth (57).

The discovery that NAD^+^, one of the most important enzyme cofactors existing in all living organisms, is attached to RNA in a large variety of organisms, including Archaea, reflects its fundamental role in nature. Future identification of NAD^+^-RNAs in archaeal organisms will determine whether archaeal noncoding sRNAs, which may be involved in rRNA modification and antisense regulation (58–60), also carry NAD^+^ caps, as it happens in non-coding RNAs of human, plant and bacterial cells (3, 8, 13). Additionally, several archaeal species, including the model organism used in this work, are capable of digesting organic compounds for methane production, which is of industrial and biotechnological interest for biofuel production and bioremediation (19, 20). Therefore, it is tempting to speculate that NAD^+^-capping occurring preferentially in transcripts involved in redox processes in many species (4, 6, 14, 57) may lead to its potential utilization in biotechnological applications. In plant and mammalian cells, it has been shown that the NAD^+^ capping landscape is affected by exposure to stress (10, 14). Identifying the NAD^+^-capped transcripts in other archaeal organisms capable of surviving and growing in extreme environments may help to further advance our understanding of the biological function of this modification, and its relation to environmental cues and stress (61, 62).

The data obtained by NAD captureSeq have revealed that mycobacterial NAD^+^-RNAs mainly consist of mRNAs whose products are involved in cell membrane and redox processes. This is reminiscent of the findings in *E. coli, B. subtilis,* yeast and mammalian cells, where NAD^+^-mRNAs code for polypeptides that function in bacterial and mitochondrial membranes (4, 6, 14, 57), reflecting the plasma membrane localization of oxidative phosphorylation in bacteria, and mitochondrial membrane in eukaryotes. Although it is not clear yet whether the NAD^+^ caps of mycobacterial mRNAs have a function, it is possible that the NAD^+^ caps tag mRNAs for transport to cellular locations where their translation products will be used. On the other hand, the appearance of NAD^+^-capped transcripts might be a consequence of transcription of these RNAs occurring in NAD^+^-rich cellular locations.

The comparison of the results of the NAD captureSeq enrichment and the intracellular amounts of the captured transcripts showed that the transcripts were detected by NAD captureSeq not because of their high expression levels. However, it is possible that the most rare NAD^+^-capped transcripts were lost during the NAD captureSeq procedure, which includes steps potentially decreasing the integrity of RNA. This is consistent with the low amount of reads in the NAD captureSeq experiment. While this work was in preparation, identification of NAD+-capped transcripts by using copper-free methods to reduce the RNA damage were published (52, 57). Future NAD^+^-RNA identification in *M. smegmatis* employing these new protocols may unveil other rare NAD^+^ capped transcripts that were not detected in our study.

Ms1 is one of the few characterized sRNAs in mycobacteria, including *M. tuberculosis* (MTS2823). It is the most abundant non-rRNA transcript present in *M. smegmatis* cells in stationary phase. This sRNA was shown to interact with the RNAP core and to regulate its intracellular level (23, 48). We have shown that the level of NAD^+^ capping of Ms1 is higher in stationary phase. This may contribute to the markedly higher biological stability of Ms1 in stationary phase compared to exponential phase (23, 48), and is consistent with the increased stability of NAD^+^-capped RNA in *E. coli* (15). In mycobacteria, the cap may provide protection against 5’ to 3’ exonucleases, such as RNase J1 (63, 64). In addition, we have shown that mycobacterial RNAP can incorporate NAD^+^ as the initiating dinucleotide into Ms1 *in vitro* with ~40% efficiency at equal concentrations of NAD^+^ and ATP, which reflects the percentage of NAD^+^-capped Ms1 detected *in vivo* (~30-50 %) in stationary phase. This is consistent with approximately equimolar levels of ATP and NAD in stressed bacteria (65). These findings define Ms1 as an attractive model transcript to study the mechanism and role of NAD^+^ capping in Actinobacteria where Ms1 is widely conserved even in evolutionarily distant species (Vaňková Hausnerová *et al.,* 2021 manuscript in preparation).

The NAD^+^-capped transcripts in *M. smegmatis* also include three validated but yet uncharacterized sRNAs IGR-2, IGR-5 and IGR-11. This finding is consistent with the frequent presence of the NAD^+^cap at the 5’ends of small non-coding RNAs in bacterial, plant and mammal cells (3, 8, 13). Interestingly, *M. smegmatis* IGR-2 is highly expressed in stationary phase phase (Figure 2B), similarly to Ms1, thereby suggesting its important regulatory role under these physiological conditions. NAD^+^ caps have also been detected in *Streptomyces,* a different Actinobacterial genus, but the exact NAD^+^-capped transcripts are unknown, and it will be of interest to identify whether sRNA in this genus carry this modification (66)

Our search for promoters from which transcription initiated with NAD^+^ in *M. smegmatis* revealed conserved motifs corresponding to −35 and −10 promoter elements recognized by sigma factors σ^A^ and σ^F^. This is consistent with the fact that the Ms1 promoter can be transcribed *in vitro* by the RNAP holoenzyme containing σ^A^ (Figure 4), and that it was previously observed that Ms1 accumulated after σ^A^ overexpression (23). σ^F^ was previously reported to be active under oxidative stress, heat shock, low pH and in stationary phase (67–69) and is also highly expressed in exponential phase (70). This is consistent with the finding that most of the NAD+ capped transcripts were detected in stationary phase. A comparison of the *E. coli* NAD^+^ incorporation favouring sequence (TSS and its surrounding sequence) with that one derived from the mycobacterial NAD^+^-capped transcripts then revealed no similarity (Supplementary Figure 6) (15, 53). Interestingly, −1C, which decreases NAD^+^-capping efficiency in *E. coli* and *B. subtilis* (4, 15), can be found in two mycobacterial NAD^+^-capped transcripts, most notably in Ms1. This suggests that mycobacteria employ different rules dictating efficient NAD^+^ incorporaton.

NAD^+^ cap removing enzymes have been identified in bacteria, mammals, yeast and plants (4, 7, 13, 47, 71–73). They are often represented by members of the Nudix family (3, 14, 74). In *M. smegmatis* and *M. barkeri,* putative NAD^+^ diphosphatases are annotated (MSMEG_1946 and Mbar_A2829, accession numbers A0QTS4 and Q468G3, respectively) but have not been experimentally validated, and future experiments are required to address NAD^+^ decapping in these organisms.

In conclusion, our data indicate that NAD^+^ capping is conserved among phylogenetically distant biological species and widespread in all kingdoms of life. Interestingly, a comparison of NAD^+^ capping in eukaryotic and prokaryotic organisms reveals a common pattern: (i) common functions of proteins encoded by NAD^+^-capped RNAs, (ii) presence of NAD^+^ caps in sRNAs, and (iii) changes in its incidence in different growth phases. Further exploration of NAD^+^ capping in other model organisms, including Archaea, will help determine the key mechanisms controlling NAD^+^ capping and its biological functions in evolutionarily distant organisms.

## AVAILABILITY

The complete bioinformatic workflow for analysis of NAD captureSeq data is available at https://github.com/mprevorovsky/krasny-NADcaptureSeq.

## SUPPLEMENTARY DATA

Supplementary Data are available at NAR Online.

## ACCESSION NUMBERS

The raw sequence read files were deposited in the ArrayExpress database (accession number E-MTAB-10246).

## ACKNOWLEDGEMENTS

We thank Viola Vaňková Hausnerová, Petra Sudzinová, and other members of the Krásný lab for their help and discussion.

## SUPPLEMENTARY MATERIAL

**Supplementary Table 1.** Enriched loci at the 5’ end of 25 genes by NAD captureSeq. The average fold enrichment between “ADPRC+” and “ADPRC-” samples in exponential and stationary phase is indicated (“Inf” indicates that the value of the signal for the “ADPRC-” sample equalled zero). All TSS were taken from Li et al., 2017 (1), except for Ms1, which was taken from Hnilicova et al., 2014 (2), and IGR-2, IGR-5 and IGR-11, that were taken from Li et al., 2013 (3). The correlations between TSS and NAD captureSeq signal for IGR-2, IGR-5 and IGR-11 are discussed in the Results section.

| NAD<sup>+</sup>-RNA candidates | Enrichment |  | Annotation | ORF 5'-end | TSS | NAD captureSeq signal |
| --- | --- | --- | --- | --- | --- | --- |
|  | Exp. phase | Stat. Phase |  |  |  |  |
| MSMEG_0944 | 18.9 | - | DNA binding domain-containing protein | 1024188 | 1024110 | 1024112 (A) |
| MSMEG_6345 | 6.2 | 11.6 | Hypothetical protein | 6407908 | 6407838 | 6407839 (A) |
| MSMEG_0266 | - | Inf | Arginine decarboxylase | 299314 | 299314 | 299314 (A) |
| MSMEG_0670 | - | Inf | FAD dependent oxidoreductase | 752392 | 752392 | 752392 (A) |
| MSMEG_1919 | - | Inf | WhiB family transcriptional regulator, redox-sensing transcriptional regulator | 2000397 | 2000588 | 2000588 (A) |
| MSMEG_4738 | - | Inf | Hypothetical protein | 4838569 | 4838415 | 4838415 (A) |
| MSMEG_6935 | - | Inf | N-acetylmuramoyl-L-alanine amidase | 6978405 | 6978348 | 6978348 (A) |
| MSMEG_6676 | - | 356.5 | Transmembrane protein | 6728933 | 6728779 | 6728779 (A) |
| MSMEG_1112 | - | 135.5 | Aconitate hydratase | 1178451 | 1178451 | 1178451 (A) |
| MSMEG_1788 | - | 43.9 | Hypothetical protein | 1871258 | 1871277 | 1871279 (A) |
| MSMEG_0987 | - | 41.8 | Hypothetical protein | 1059955 | 1059900 | 1059900 (A) |
| MSMEG_3689 | - | 33.5 | Sodium:solute symporter | 3754973 | 3755061 | 3755061 (A) |
| MSMEG_2756 | - | 32.0 | Hypothetical protein | 2825374 | 2825424 | 2825424 (A) |
| MSMEG_2010 | - | 30.8 | Stress responsive A/B barrel domain superfamily protein | 2089956 | 2090033 | 2090035 (A) |
| MSMEG_5081 | - | 29.4 | Hypothetical protein | 5181146 | 5181247 | 5181247 (A) |
| MSMEG_5617 | - | 19.1 | Immunogenic protein MPT63 | 5701302 | 5701364 | 5701364 (A) |
| MSMEG_5246 | - | 14.0 | Hypothetical protein | 5338643 | 5338643 | 5338643 (A) |
| MSMEG_1605 | - | 10.8 | Phosphate transporter regulatory protein PhoU | 1692222 | 1692182 | 1692182 (A) |
| MSMEG_5648 | - | 9.6 | Hypothetical protein | 5737325 | 5737325 | 5737325 (A) |
| MSMEG_1519 | - | 8.9 | Translation initiation factor IF-1 | 1611597 | 1611453 | 1611453 (A) |
| MSMEG_4268 | - | 5.0 | Cytochrome C oxidase subunit 2 | 4352234 | 4352293 | 4352293 (A) |
| Ms1 | - | 3.9 | Regulatory small RNA | - | 6242368 | 6242368 (A) |
| IGR-2 | - | 7.6 | Putative small RNA | - | 417847 | 417879 (A) |
| IGR-5 | 5.9 | Inf | Putative small RNA | - | 5862669 | 5862639 (A) |
| IGR-11 | 3.9 | 31.1 | Putative small RNA | - | 396650 | 396831 (A) |

**Supplementary Table 2:**
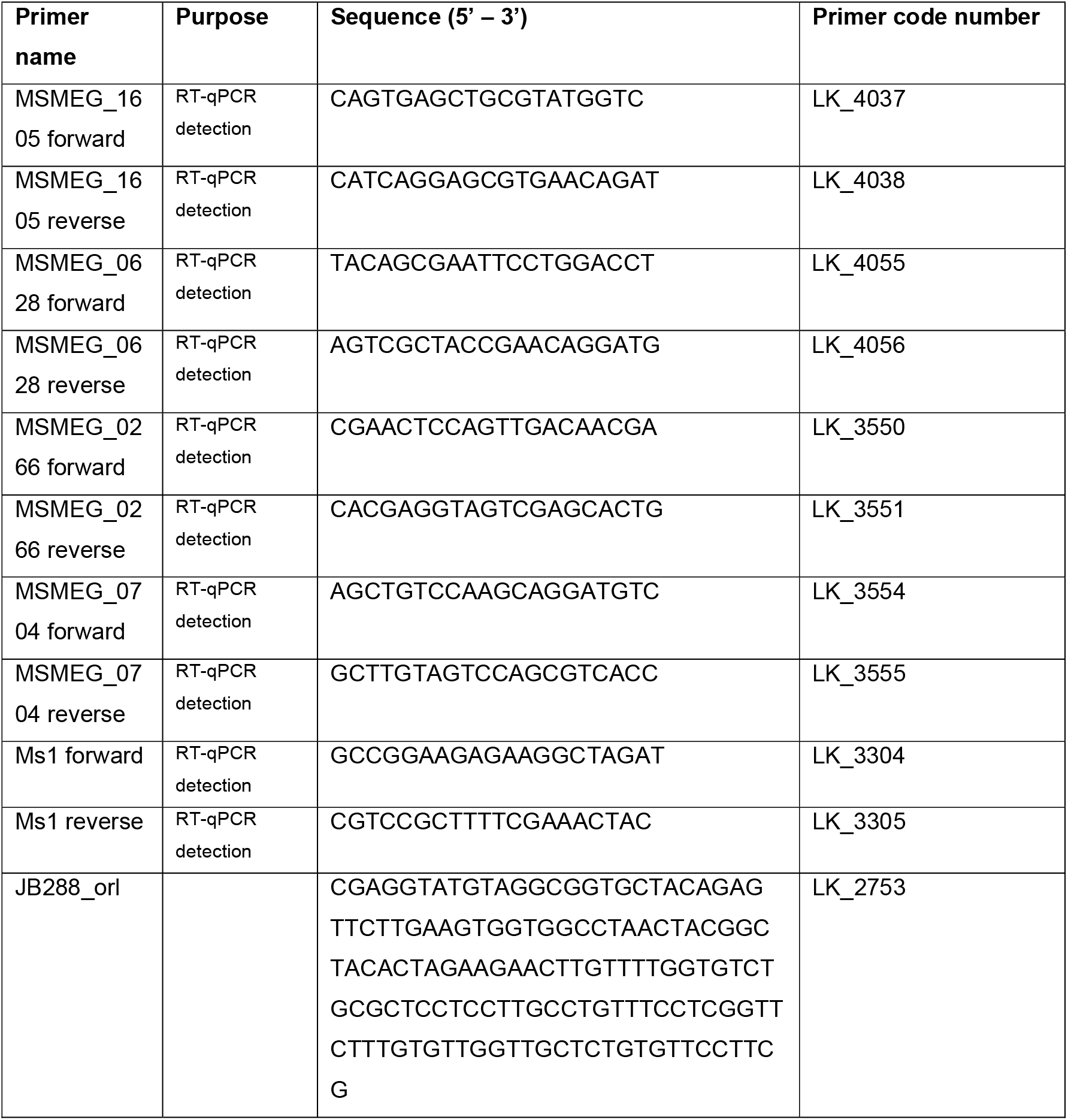
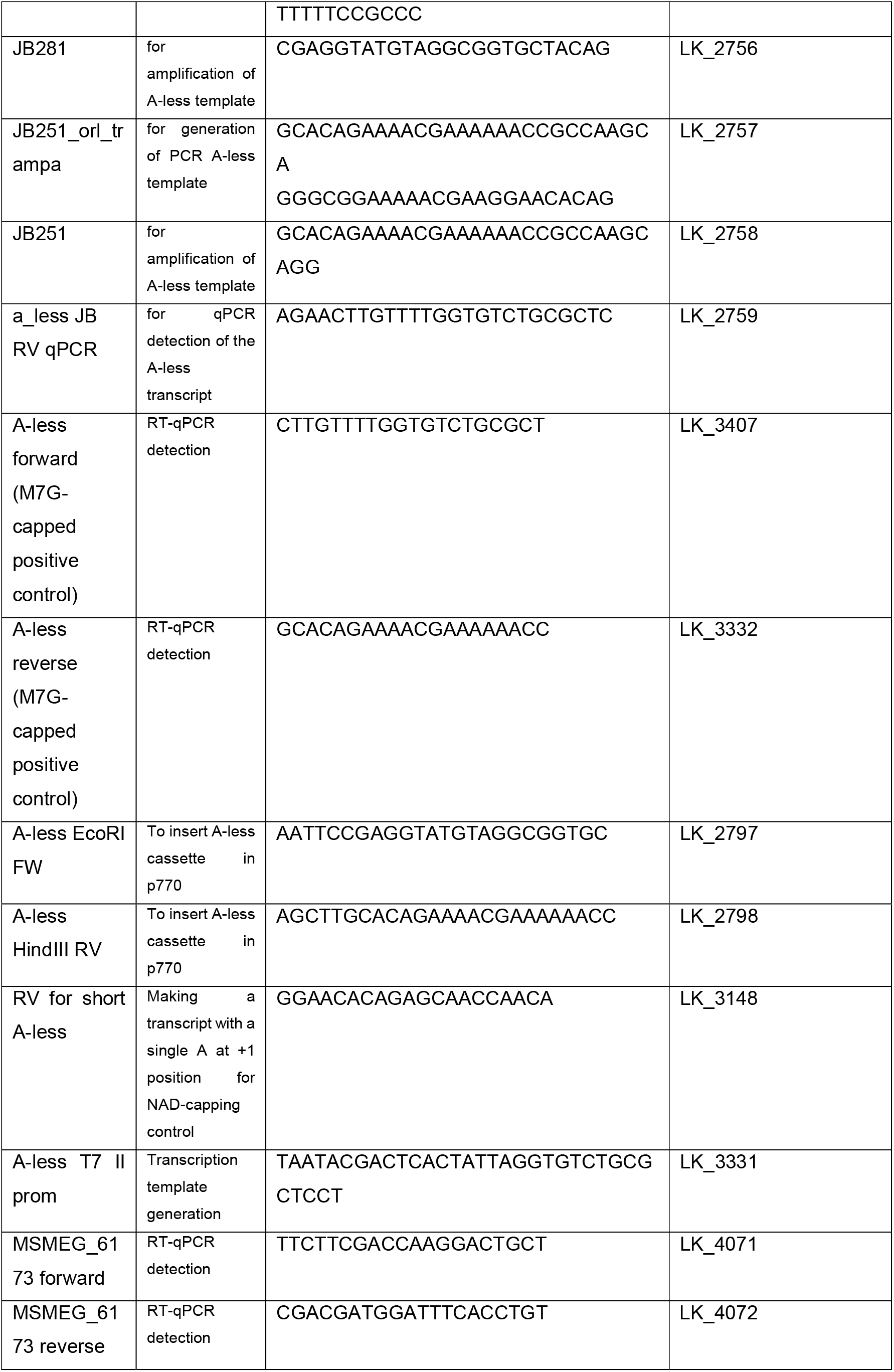
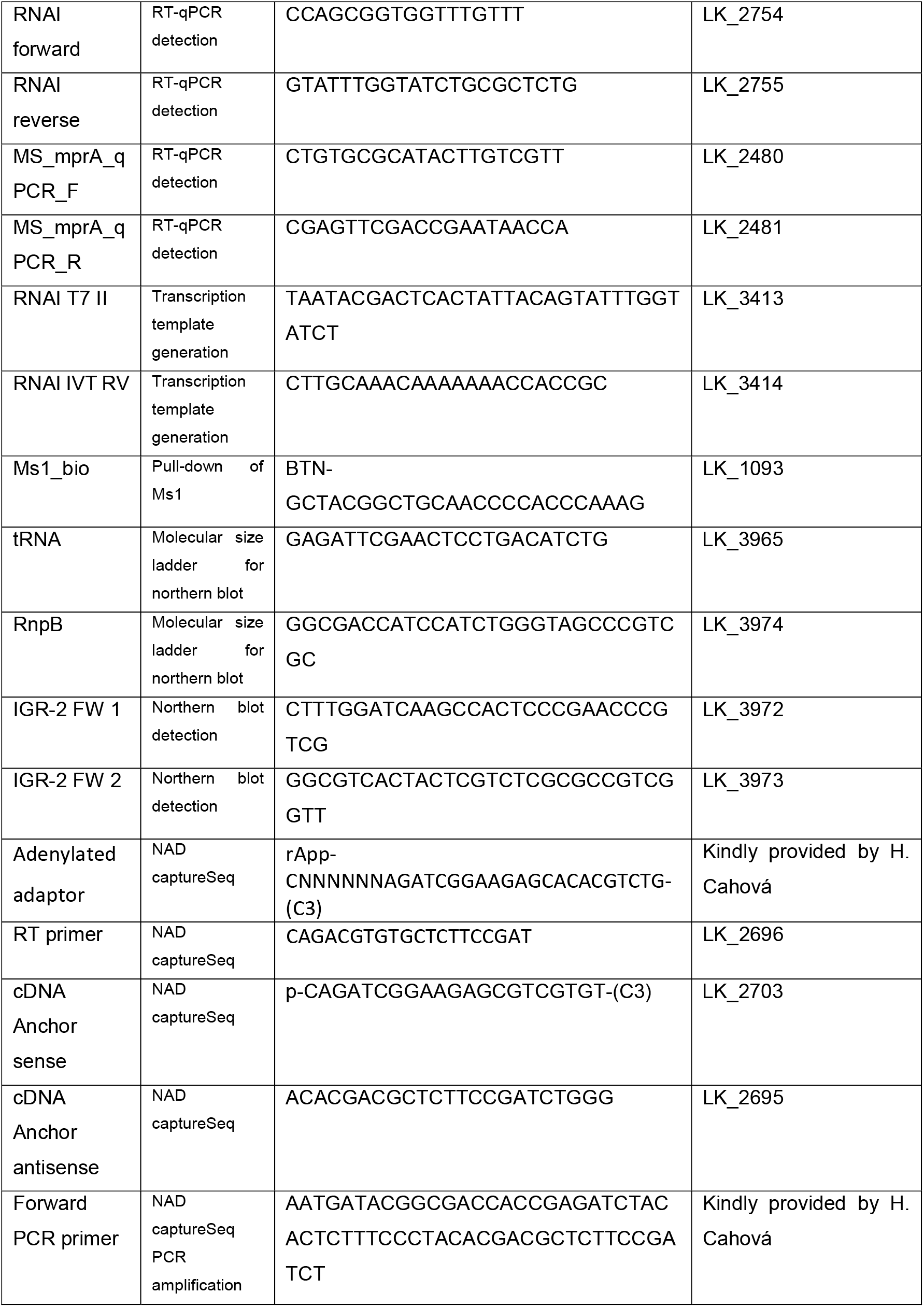
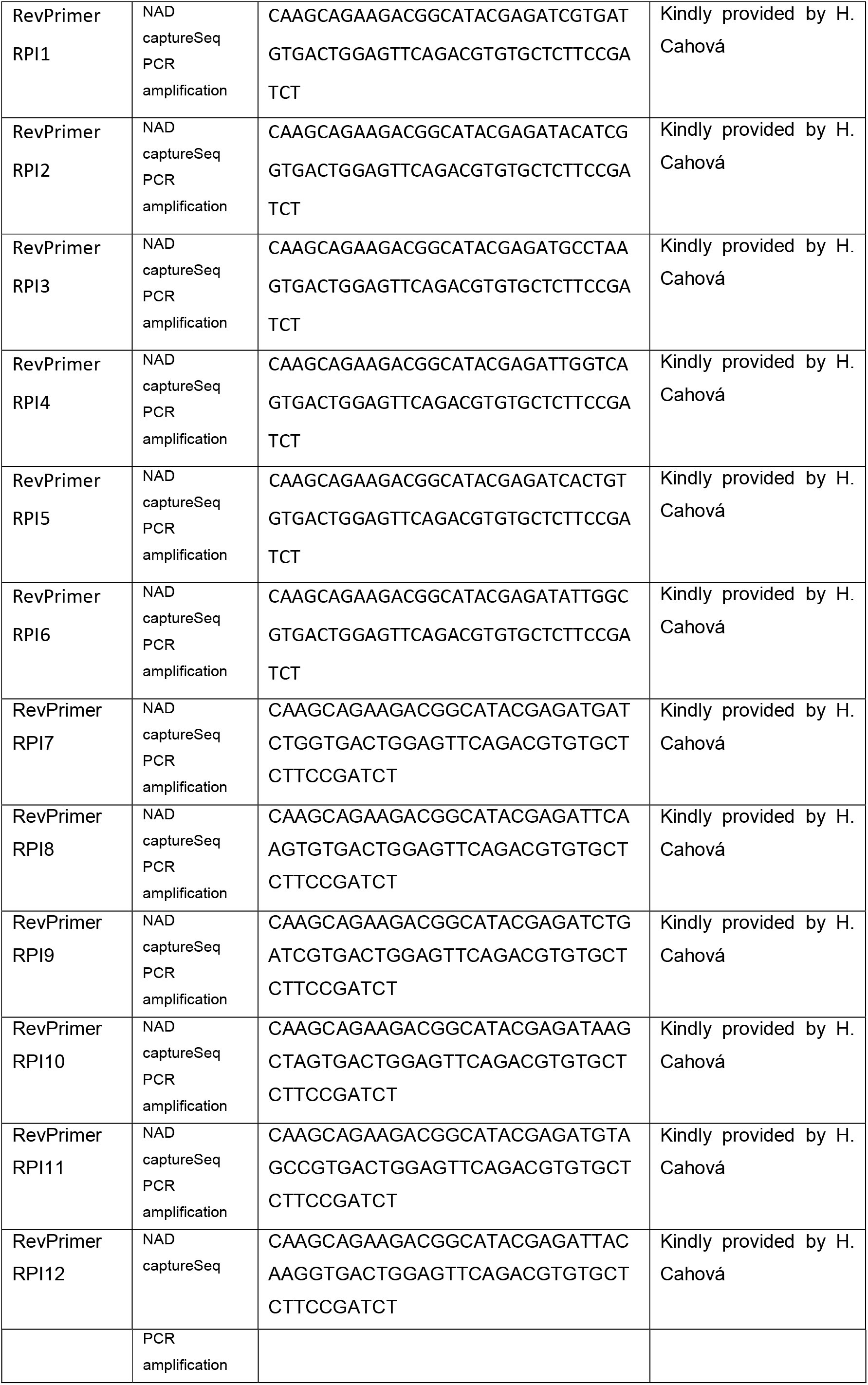
Primers used in this work.

**Supplementary Figure 1.**
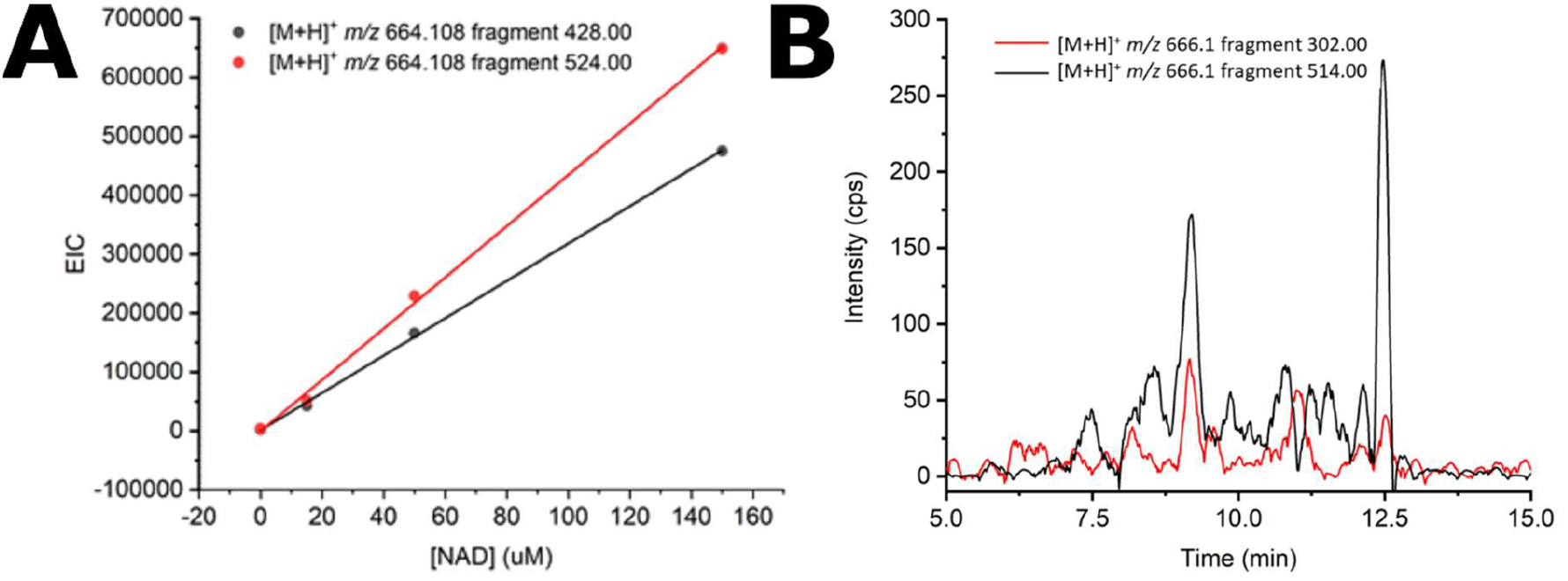
**A.** Example of regression curves used for absolute quantification of NAD^+^ from total RNA of *M. smegmatis* from stationary phase. 10 μl of 0 μM, 15 μM (150 pmol), 50 μM (500 pmol) and 150 μM (1.5 nmol) standard NAD^+^ solution were added to 500 μg aliquots of RNA and analysed by MRM. The intersection of the lines created by detection of the two different fragments of NAD^+^ ([M+H]+ 428.00 and 524.00) equals the concentration of NAD^+^ in the sample. **B.** MRM Chromatogram of NADH from RNA of M. *smegmatis* in stationary phase where two different fragments ([M+H]^+^ 302.00 and 514.00) were detected.

**Supplementary Figure 2.**
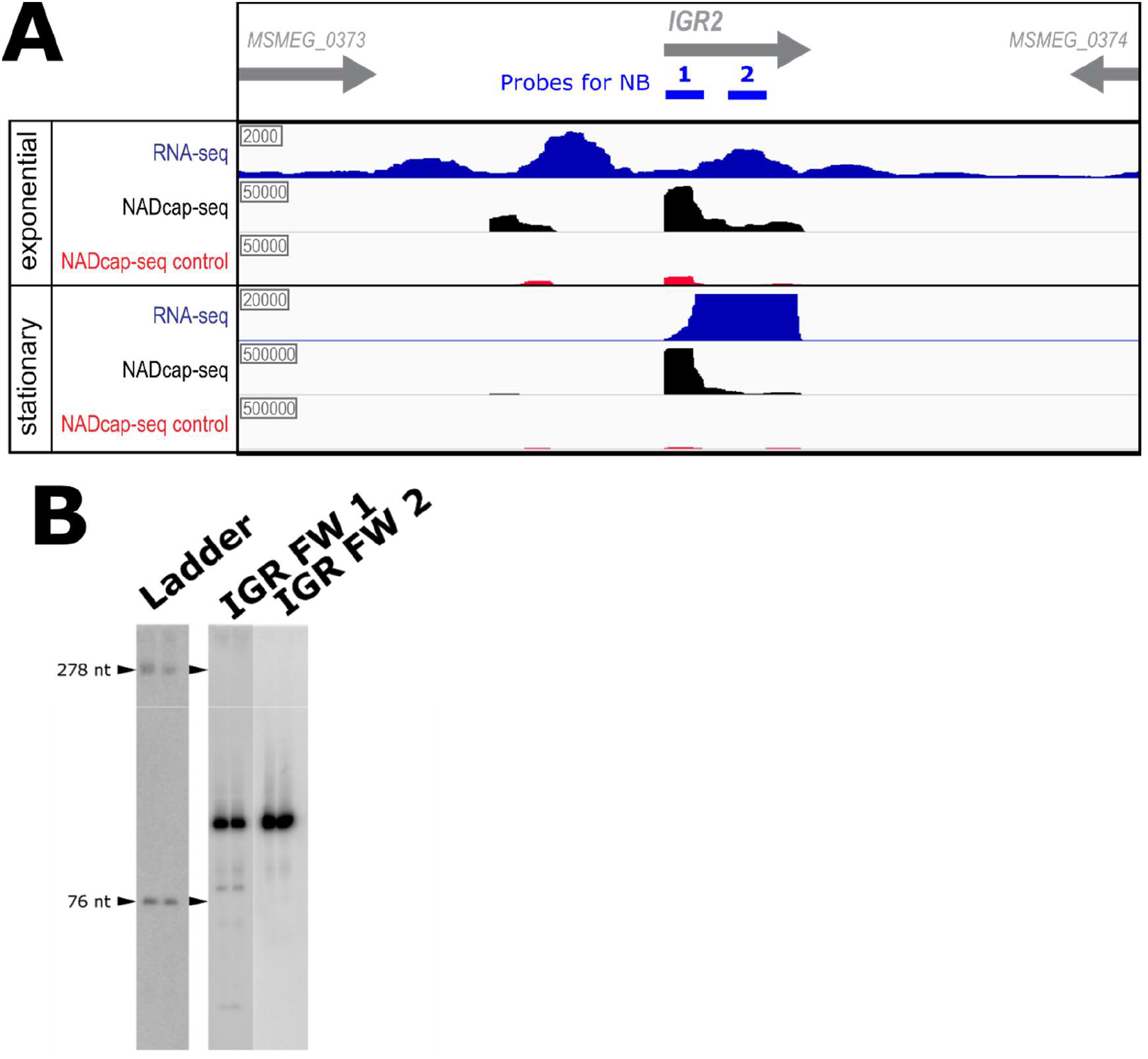
Northern blot analysis of IGR-2. **A.** NAD CaptureSeq (incl. negative control) and RNAseq data for the MSMEG_0373-MSMEG_0374 intergenic region. The signal detected by a previous RNA-Seq study is in blue (48), the signal corresponding to the NAD captureSeq in black, and the signal corresponding to the non-biotinylated negative control in red. The probes (1,2: IGR-2 FW 1 and IGR-2 FW 2) used for Northern blot analysis of IGR-2 are shown in blue in the upper part of the pannel. **B.** Northern blot results show at least 2 species of IGR-2 with approximate sizes of 83 and 123 nucleotides long (more prominent), respectively. The approximate lengths were estimated using GelAnalyzer software, http://www.gelanalyzer.com/). Each probe was used on membranes with transferred biological duplicates of total RNA samples extracted from *M. smegmatis* cells in stationary phase.

**Supplementary Figure 3.**
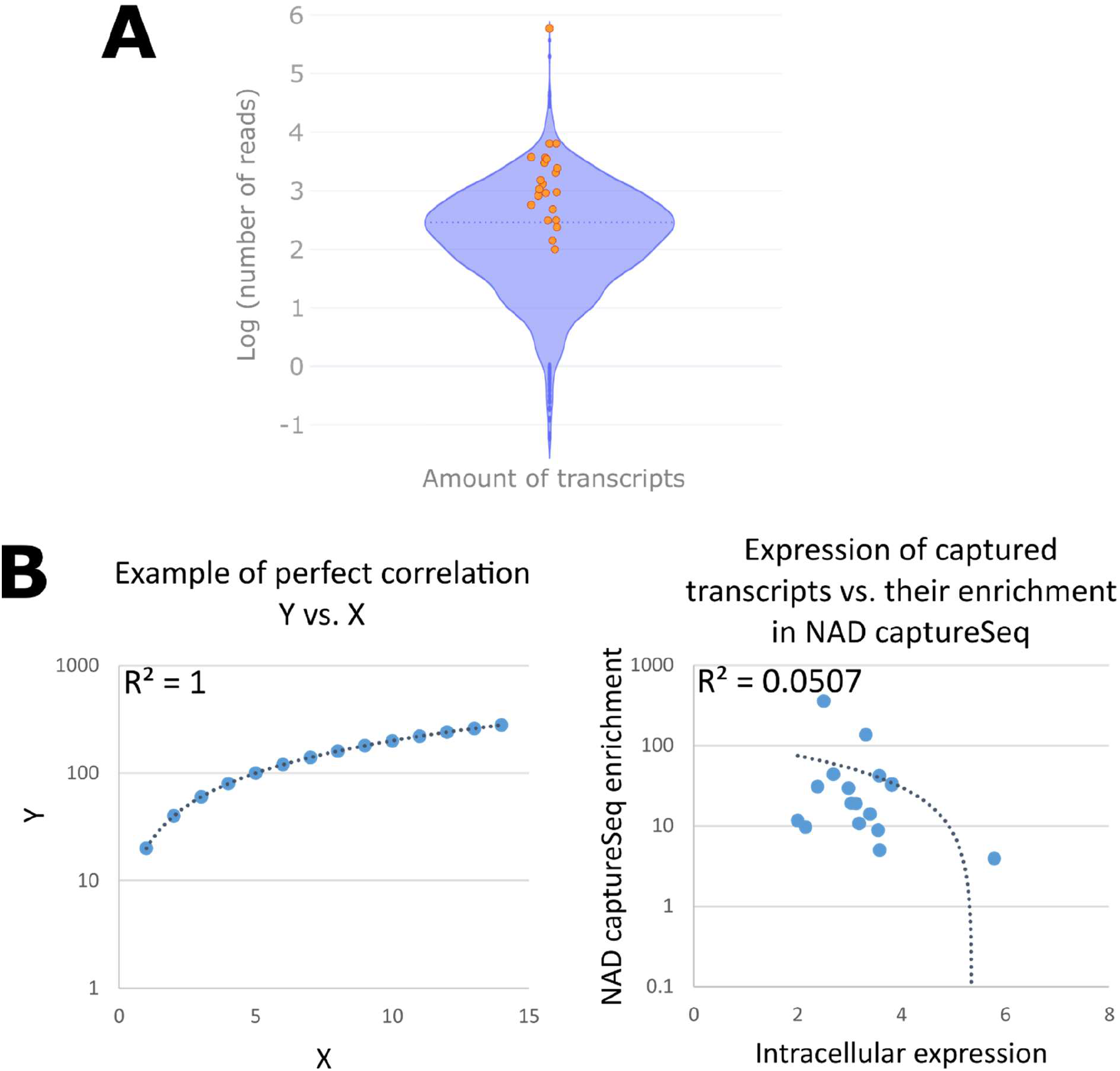
The enrichment detected by NAD captureSeq experiment is not caused by the high abundance of the transcripts. **A.** The logarithmic values of normalised number of reads of 22 transcripts detected by NAD captureSeq (orange dots) are plotted on the expression of all transcripts detected by RNA-Seq (blue violin plot) (4) (IGR-2, IGR-5 and IGR-11 were not annotated among the detected transcripts by RNA-seq experiment and were excluded from the plot). The violin plot shows the distribution of the expression of the transcripts. The position of transcripts detected by NAD captureSeq is moderately over the median expression of all transcripts *in M. smegmatis* (dashed line). The highest point represents the expression level of Ms1 sRNA. The violin plot was made using Plotify software (chart-studio.plotly.com), Microsoft Excel and Inkscape. **B.** Side-by-side comparison of graphs representing a perfect direct proportionality between variables (left) and the representation of the expression of the captured transcripts as detected by RNA-Seq (4) and their enrichment in the NAD captureSeq experiment (right). The comparison of both graphs shows that there is no apparent correlation between the abundance of the transcripts and their enrichment in NAD captureSeq.

**Supplementary Figure 4.**
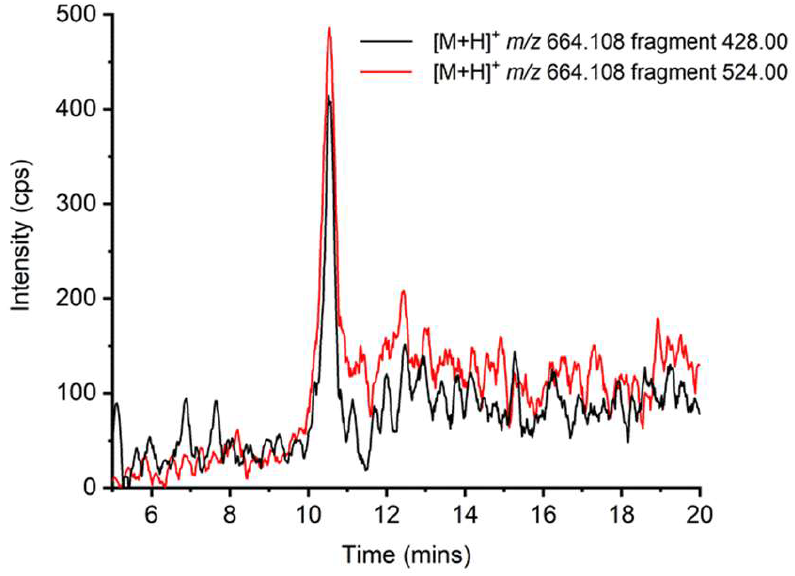
Validation of Ms1 NAD^+^ capping by Mass Spectrometry. MRM Chromatogram of NAD^+^ from Ms1 of M. *smegmatis* in stationary phase where two different fragments of NAD^+^ were detected ([M+H]^+^ 428.00 and 524.00, see Figure 1D).

**Supplementary Figure 5.**
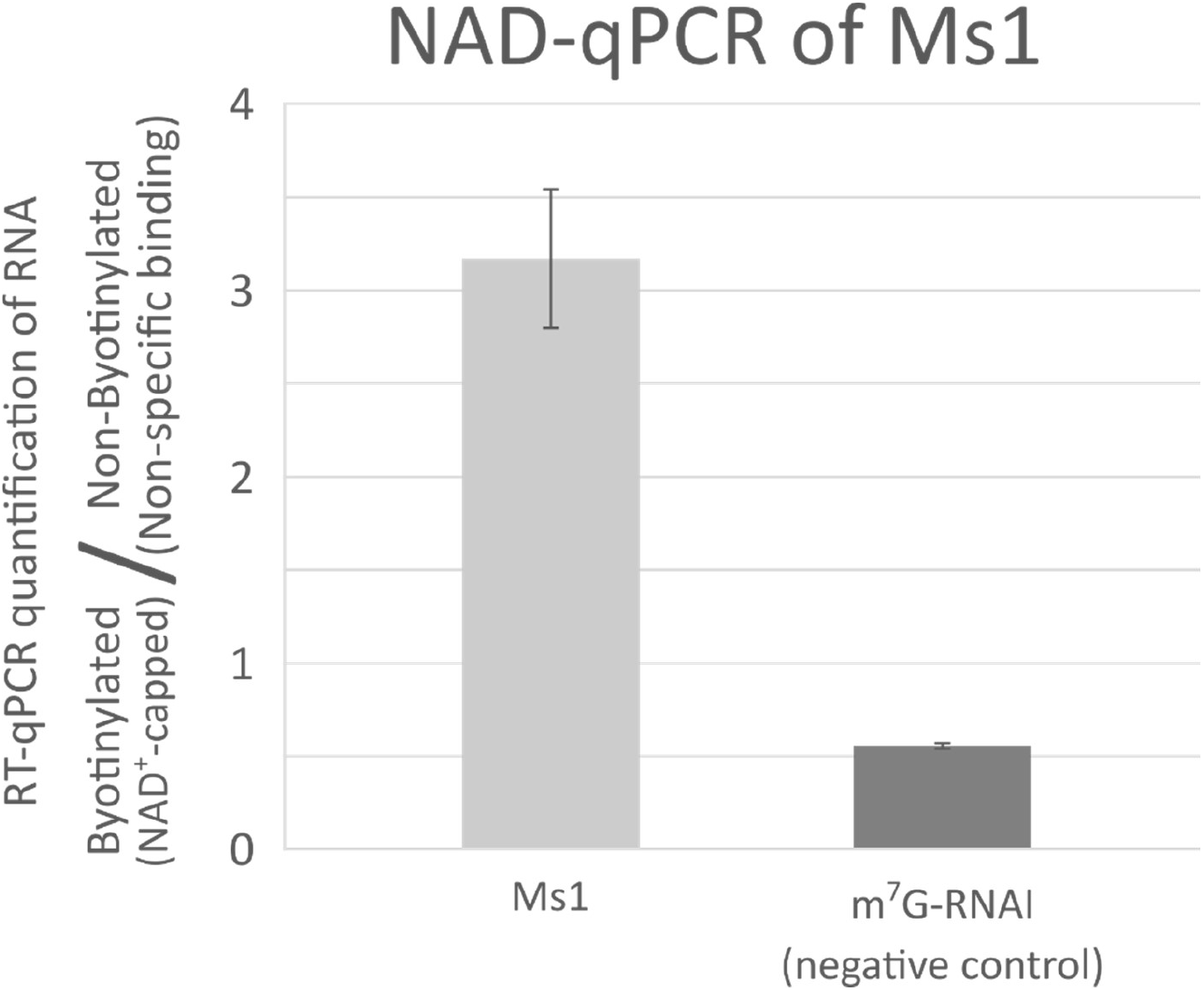
Validation of Ms1 NAD^+^ capping by NAD-qPCR. Total RNA from M. *smegmatis* was spiked with an m^7^G-capped transcript as a negative control. NAD^+^ at the 5’ end of RNA was substituted with biotin for specific capture of the NAD^+^-capped RNA (Biotinylation) or mock treated for detection of non-specific binding to the streptavidin beads (Non-Biotinylation). The aliquots of RNA were added unto streptavidin beads, and the amount of Ms1 and the negative control captured was evaluated by RT-qPCR. The plot shows ratios between detected Ms1 and negative control in the biotinylated aliquot (indicating specific capture of NAD^+^-capped transcripts) and biotinylated (or mock treated) aliquot (indicating detection of non-specifically bound transcripts). The bars show averages from two experiments and the error bars show the range ±SD.

**Supplementary Figure 6.**
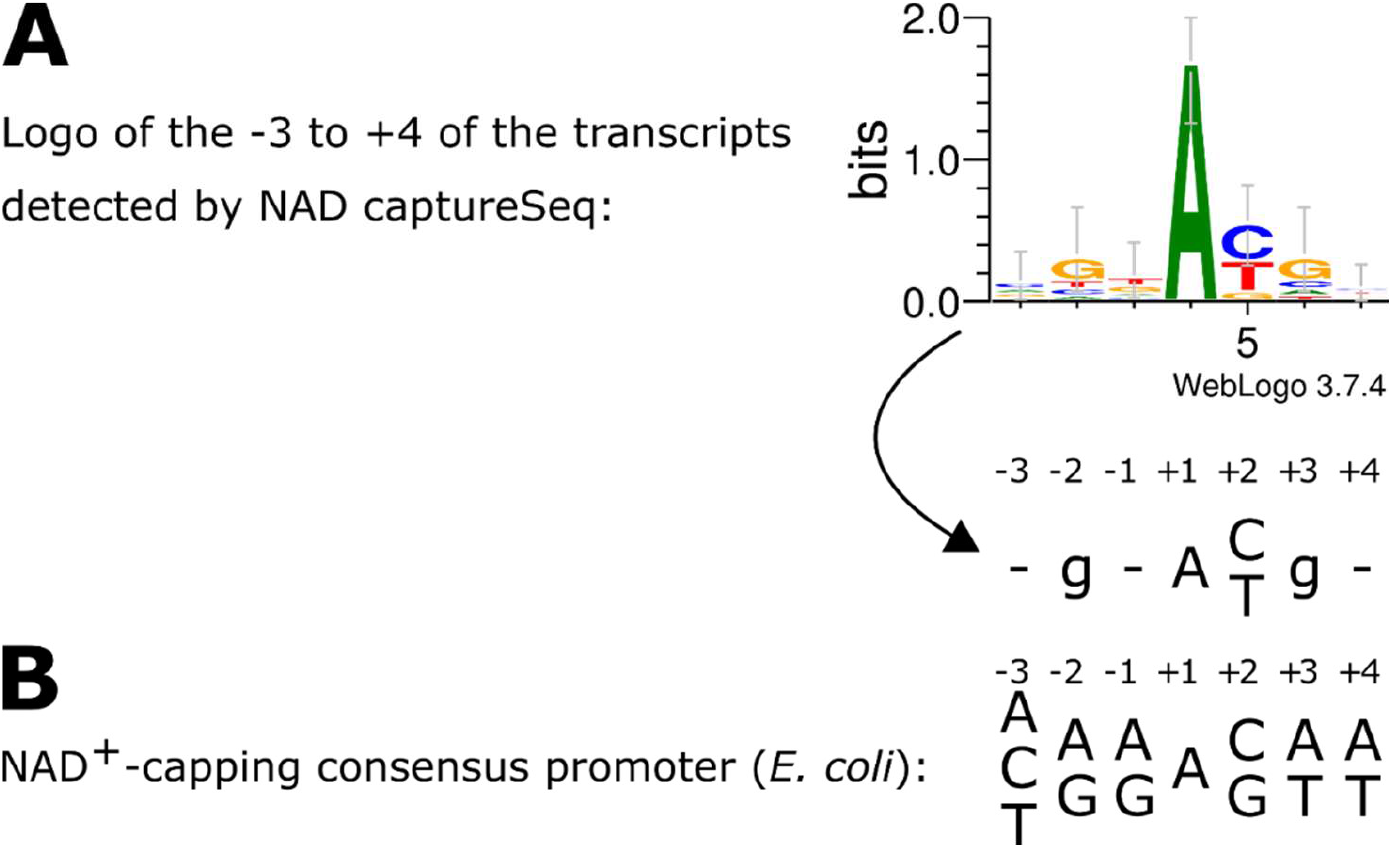
The −3 to +4 logo of NAD^+^-capped transcripts from *M. smegmatis* does not resemble the consensus promoter for NAD^+^ capping in *E. coli*. A. The −3 to +4 sequence of all NAD captureSeq detected genes were used to determine their sequence logo using the WebLogo software (http://weblogo.threeplusone.com). **B**. Previously defined consensus promoter for NAD^+^-capping of *E. coli* (6).

